# Cortical Stimulation Strength and sgACC Connectivity Shape Neuro-Cardiac Responses to Prefrontal TMS

**DOI:** 10.64898/2026.05.10.724073

**Authors:** Zijian Feng, Sandra Martin, Gerasimos Gerardos, Konstantin Weise, Gesa Hartwigsen, Thomas Knösche, Ole Numssen

## Abstract

Autonomic dysregulation is associated with major depressive disorder, yet how cortical stimulation strength and network connectivity shape autonomic responses to prefrontal transcranial magnetic stimulation (TMS) remains unclear. We combined resting-state functional MRI, individualized electric-field simulations, and electrocardiography during repetitive TMS at up to six left dorsolateral prefrontal cortex (DLPFC) targets in 15 healthy adults. Heart–brain coupling (HBC), quantified as cardiac spectral power at the stimulation-cycle frequency, was modeled using either motor-threshold-based intensity or cortical electric-field strength, with and without DLPFC connectivity to the subgenual anterior cingulate cortex (sgACC). In linear mixed-effects models, electric-field strength provided better fit and explained more HBC variance than motor-threshold-based intensity. Adding sgACC connectivity further improved model fit and revealed site-specific electric-field-by-connectivity interactions. High-resolution mapping revealed spatial correspondence between regions of higher E-field–HBC explained variance and sgACC-anticorrelated cortex. Negative sgACC connectivity was associated with steeper predicted increases in HBC at several sites, while nonlinear models showed site-dependent differences in dose–response profiles between negative and positive connectivity. Together, these findings link acute cardiac responses to cortical electric-field exposure and sgACC connectivity, supporting HBC as a network-sensitive physiological readout for investigating frontal–vagal function. This framework provides a basis for testing individualized prefrontal TMS targeting and dosing in depression-related autonomic dysregulation.

## Introduction

Major depressive disorder (MDD) is accompanied by autonomic dysregulation, including elevated resting heart rate and reduced heart rate variability, reflecting diminished cardiac vagal modulation [1, 2]. The prefrontal cortex plays a central role in cardiovascular autonomic control [3]. Recent meta-analyses confirm that non-invasive brain stimulation, particularly repetitive transcranial magnetic stimulation (rTMS) targeting the prefrontal cortex, can reduce heart rate, augment heart rate variability [4, 5], and improve outcomes in treatment-resistant depression [6, 7]. Yet, it remains unclear how the downstream connectivity of prefrontal stimulation sites shapes these autonomic effects in humans, and whether individualized, network-informed models of cortical stimulation can systematically harness them.

The observed downstream heart rate effects align with the frontal–vagal network theory, which posits that the dorsolateral prefrontal cortex (DLPFC), subgenual anterior cingulate cortex (sgACC), and vagus nerve form an integrated circuit central to both MDD pathology and autonomic regulation [8–10]. The Neuro-Cardiac-Guided TMS (NCG-TMS) framework leverages this brain–heart link by using TMS-evoked heart rate changes as real-time physiological feedback to probe frontal–vagal function: a brief on–off rTMS paradigm (“10 Hz Dash”; 5 s on / 11 s off, 0.0625 Hz) entrains the inter-beat interval to the stimulation rhythm, and the resulting heart–brain coupling (HBC) is quantified as spectral power at the TMS frequency in the cardiac signal [11]. In a recent study using the same dataset [12], we applied the NCG-TMS 2.0 protocol across three sessions in healthy participants and demonstrated robust target- and intensity-specific HBC in the DLPFC: Stimulation targets surrounding Beam F3 elicited HBC above sham, with an increase as a function of stimulator intensity expressed in percentage of individual motor threshold (%MT) (Fig. 1). However, these analyses relied on conventional stimulator-based dosing metrics and incorporated neither the TMS-induced cortical electric-field (E-field) nor the functional connectivity (FC) architecture of the stimulated tissue.

**Figure 1.**
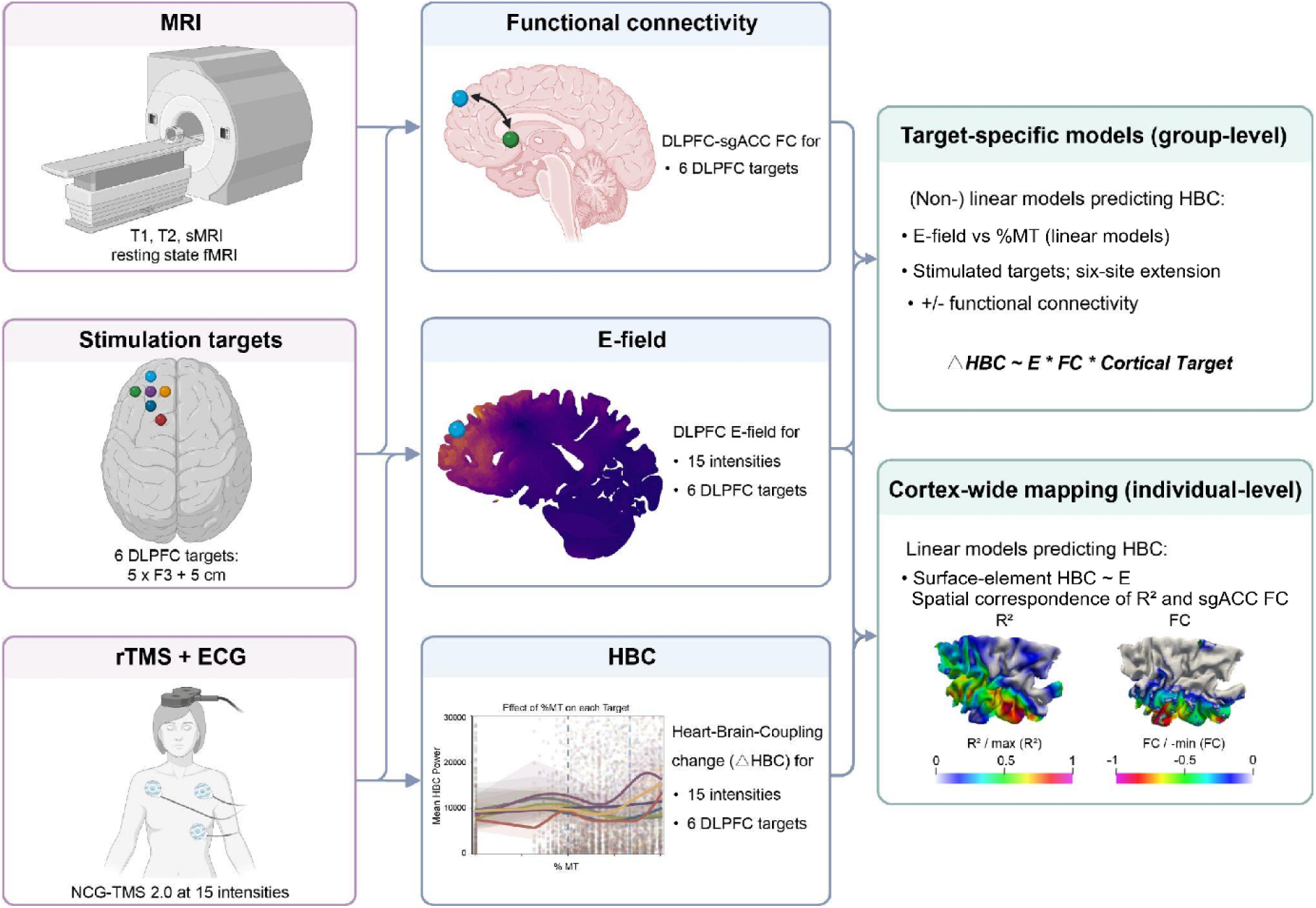
Study design. Resting-state and structural MRI were acquired to estimate individual dorsolateral prefrontal cortex (DLPFC) functional connectivity (FC) with the subgenual anterior cingulate cortex (sgACC) and to build head models for electric-field (E-field) simulations. Six left DLPFC stimulation targets were defined.

In parallel, rTMS targeting for depression increasingly relies on individual or group-level FC between DLPFC and downstream regions. Antidepressant outcomes tend to be stronger when stimulation is delivered to DLPFC sites showing stronger negative FC with the sgACC, motivating connectivity-guided targeting based on peak sgACC anticorrelation in resting-state fMRI [13–17]. A recent study showed that the association between DLPFC–sgACC FC and clinical improvement is stronger when the stimulated region is defined via E-field modeling rather than scalp coordinates [18]. In a complementary fashion, TMS-induced HBC was often strongest for individualized prefrontal targets negatively connected to the sgACC [19], positioning HBC as a potential fMRI-free readout of frontal–sgACC circuit engagement. Recent intracranial work further shows that prefrontal subregions exhibit distinct directed connectivity patterns with sgACC and large-scale networks, underscoring the importance of precise target selection within DLPFC [20, 21]. At the same time, individualized sgACC-based FC targeting is constrained by substantial resource demands.

Beyond identifying an optimal stimulation target [22], dosing poses an equally non-trivial challenge [23]. Standard practice calibrates TMS intensity as a percentage of individual motor threshold, implicitly assuming equivalent cortical excitability across regions and individuals. In reality, individual differences in scalp-to-cortex distance, skull thickness, and gyral geometry introduce substantial variability in the cortical E-field induced by TMS across subjects and sites [24–28]. Calibrating stimulation to individual cortical E-field rather than %MT has been proposed to reduce this variability, but implementations in clinically relevant protocols remain sparse [24, 25, 29]. Despite parallel advances in FC-guided targeting, E-field dosimetry, and neuro-cardiac TMS readouts, no study has yet combined all three within a single, individualized modeling framework.

Key to overcoming these limitations is a shift from a traditional coil-scalp-based perspective to an individualized, cortex-centered view that explicitly models the stimulated tissue, analogous to moving from sensor to source space in EEG/MEG (or from k-space to image space in MRI). In this study, we apply a cortical TMS “source-space” framework to repetitive stimulation of higher association cortex, combining individual E-field simulations for each target and intensity with DLPFC–sgACC connectivity and downstream cardiac readouts. Using this framework, we demonstrate that cortical stimulation–response mappings better explain downstream TMS effects than conventional mappings based on the motor threshold metric. By integrating DLPFC–sgACC connectivity in this cortical mapping approach, we characterize how network context relates to cortical E-field–HBC relationships.

## Materials and methods

### Participants and study design

TMS and HBC data were reused from a previously published, pre-registered study that examined target specificity and test–retest repeatability of DLPFC stimulation sites on HBC [12]. For the present analyses, we included the subset of participants (15 out of 19 healthy adults; 7 females; age 18–39 years, mean ± SD 31.6 ± 5.7 years) for whom the required MRI data were available, enabling FC estimation, pulsewise coil displacement (PCD) quantification, and individual E-field simulations. All participants were right-handed according to the Edinburgh Handedness Inventory [30] and underwent medical screening for TMS eligibility following established safety guidelines [31]. The study complied with the Declaration of Helsinki and was approved by the Ethics Committee of the Medical Faculty at Leipzig University (342/23-ek); all participants provided written informed consent and received financial compensation. Full details on recruitment and exclusion criteria are reported in [12].

### Data acquisition and experimental design

#### MRI

For each participant, MRI scans were acquired on a 3T scanner (Siemens Verio or Skyra fit) with a 32-channel head coil. Structural T1- and T2-weighted images were obtained for neuronavigation, E-field simulation, and as spatial reference for resting-state scans. T1-weighted images (MPRAGE ADNI) were acquired with 176 sagittal slices, repetition time (TR) = 2.3 s, echo time (TE) = 2.98 ms, field of view (FoV) = 256 mm, voxel size = 1 × 1 × 1 mm³, no slice gap, and flip angle = 9°. T2-weighted images were acquired with 192 sagittal slices, matrix size = 256 × 258, voxel size = 0.488 × 0.488 × 1 mm³, flip angle = 120°, and TR/TE = 5000/395 ms. If high-quality structural scans (< 2 years old) were available in the in-house database, they were used instead of new acquisitions.

Resting-state fMRI data were acquired on a 3T Siemens Prisma fit with a 64-channel head coil using a multiband (MB = 6) gradient-echo EPI BOLD sequence (TR = 1.0 s, TE = 30.8 ms, flip angle = 60°, FoV = 220 mm, voxel size = 2 mm isotropic). Phase encoding was anterior–posterior (AP) with a pixel bandwidth of 2165 Hz/Px. The fMRI lasted ∼10 minutes, with the first 10 seconds (10 volumes) discarded. A single-volume reverse phase-encoded EPI (posterior–anterior, PA) with identical geometric parameters was acquired for susceptibility-induced distortion correction.

#### TMS

TMS was delivered using a MagPro X100 stimulator (MagVenture, Farum, Denmark) with a passively cooled MCF-B65 figure-of-eight coil. Coil positioning was guided by a neuronavigation system (TMS Navigator, Localite, Sankt Augustin, Germany) with a Polaris Spectra camera (NDI, Waterloo, Canada). At the beginning of the first session, the individual motor threshold (MT) for the first dorsal interosseous (FDI) muscle was determined following published guidelines [32]. Six targets within the left DLPFC were defined: Beam F3, F3 anterior, F3 posterior, F3 lateral, F3 medial (each 2 cm from F3), and a 5-cm rule target. The 5-cm condition was acquired in eight of the 15 MRI participants; only recorded stimulation conditions were analysed. The coil was oriented at 45° relative to the parasagittal plane (handle pointing posteriorly), with adjustments to 90° for participants reporting pain scores > 6 on the Defense and Veterans Pain Rating Scale (DVPRS). The order of coil placements was randomized and kept constant within subjects across the three sessions. Participants were blinded to stimulation conditions.

The NCG-TMS 2.0 protocol [11] consisted of 16 blocks per target: one rest block [0 % *maximum stimulator output* (%MSO)] followed by 15 active blocks with intensity increasing in step sizes of 2% MSO up to 120% MT. Each block comprised a 5 s 10 Hz TMS train followed by an 11 s inter-trial interval. Each target assessment lasted approximately 7 min, including pre- and post-stimulation resting ECG recordings, as described previously [12]. The main data acquisition (16 blocks x 16 s each) thus takes about 4.2 minutes per target. After each session, participants completed a standardized side-effects questionnaire [33].

#### ECG

ECG data were recorded during each target stimulation using a REFA8 68-channel amplifier system (TMSi, Oldenzaal, The Netherlands) and Brain Vision Recorder software (Brain Vision, MedCaT B.V.). Electrodes were positioned with the ground above the left breast, the reference above the right breast, and the active electrode below the left breast. ECG signal quality was visually inspected before each session.

### Preprocessing

#### Functional connectivity

Resting-state and T1-weighted MRIs were preprocessed with fMRIPrep (v25.1.3) [34], including head-motion, slice-time, and susceptibility-distortion correction, resampling to native T1 space, and spatial normalization to the MNI152NLin2009cAsym template. FC was computed in native T1 space using nilearn (v0.11.1) [35]. The bilateral sgACC seed comprised the union of two 10-mm-radius spheres centred at MNI (±6, 16, −10), following Sun et al. [36]. Spheres were constructed after transforming their centres to native T1 space. Seed and whole-grey-matter masks were constrained by grey-matter probability ≥0.3 and resampled to the BOLD grid using nearest-neighbour interpolation. Deconfounding included high-pass regressors, motion parameters and derivatives, white-matter/CSF signals, scrubbing and GSR [37], with framewise-displacement and standardized DVARS thresholds of 0.5 mm and 5, respectively. Time series were detrended and standardized. Cleaned, unsmoothed seed-voxel time series were averaged across the union mask. Whole-grey-matter BOLD data were spatially smoothed before cleaning, using 3-mm FWHM for the group models and 8-mm FWHM for spatial mapping. Pearson correlations with the seed time series were Fisher z-transformed. Individual FC maps were projected onto the triangulated left DLPFC surface ROI, comprising FreeSurfer-derived Brodmann areas 8, 9, 10 and 46, using corrected surface-projection code based on pyNIBS v0.2026.1 [38]. For the group models, target FC was the median of non-zero projected Fisher-z values within each 10-mm-radius target extraction sphere. Sensitivity analyses used a single right-centred sgACC seed at MNI (6, 16, −10) [13, 16, 39], with otherwise identical processing.

#### Heart–brain coupling

HBC analyses followed our previous adaptation [12] of the original NCG-TMS 2.0 pipeline [11]. ECG data from our previous study [12] had already been preprocessed to obtain R-peak time series for baseline and stimulation periods; signals were band-pass filtered (0.5–20 Hz, bidirectional fourth-order Butterworth), and R-peaks were detected using SciPy and NeuroKit2, followed by visual inspection with manual correction where the automatic R-peak detection failed. No recordings in the present sample were excluded under the original >10% manual R-peak correction criterion. Manual interpolation involved two participants and affected at most 0.96% of R peaks in any retained active recording. HBC was defined as the spectral power at 0.0625 Hz, which corresponds to the onset frequency of the TMS bursts. This frequency arises from one full TMS cycle consisting of 5 s of stimulation followed by 11 s of rest. HBC was calculated from the first burst at 0% intensity to the last burst at 120% MT. In line with our previous work [12], we used both the raw HBC power and its relative change from the start of each target, i.e. stimulation intensity 0%, defined as ΔHBC_IntX_ := log(HBC_IntX_) - log(HBC_Int0_), to enable meaningful comparisons across and within sessions and targets.

#### E-field simulation and pulse-wise coil displacement

Individual head models were constructed with the charm pipeline [40] of SimNIBS (v4.5) [41] from T1- and T2-weighted images, using FreeSurfer [42] to improve grey-matter surface quality. A high-resolution DLPFC ROI (∼30,000 elements) was defined in the cortical mid-layer to mitigate boundary effects from neighboring tissue types. Whole-head E-fields for 1 A/µs were computed for each available active TMS coil placement saved by the neuronavigation system, separately for each session to capture any session-specific coil positioning. E-fields were interpolated to the DLPFC mid-layer ROI and scaled by the realized stimulation intensities (in A/µs) of the 15 stimulation levels to reflect the E-field strength in V/m. To account for stimulation variance from coil movement, we calculated the pulsewise coil displacement (PCD) [43] for each TMS pulse relative to the first tracked coil position for that TMS condition, interpolated missing samples (181 of ∼200,000), and averaged PCD within blocks. Blocks with PCD at or above the group-level 99th percentile (4.4479) were excluded from the group-model analyses.

### Analyses

#### Comparing stimulator output strength with cortical E-field exposure

Previously, we showed that stimulation strength explained HBC for specific coil placements, with stimulation strength defined relative to each participant’s motor threshold (%MT) in % of maximum stimulator output (% MSO) [12]. This coil-based dosing ignores individual anatomical differences, such as scalp-to-cortex distance differences between the motor cortex, where MT is assessed, and the DLPFC, where TMS is applied [44]. Here, we contrasted this approach with an E-field–informed strategy by extracting the cortical E-field at six predefined DLPFC sites corresponding to the nearest cortical location under each of the six coil placements.

For each site, we computed the E-field magnitude ∣E∣ (in V/m) within a 10 mm radius sphere in the DLPFC grey matter ROI under the coil centers and used the 99th percentile as a robust proxy of the local E-field maximum. Because no consensus exists on E-field extraction parameters, we repeated the LMM analyses with an alternative extraction sphere (5 mm radius) [45] to assess robustness. We then fit two linear mixed-effects models (lme4; Bates et al. [46]), one using stimulator output (%MT) and one using cortical E-field (V/m) as the primary dosing regressor:

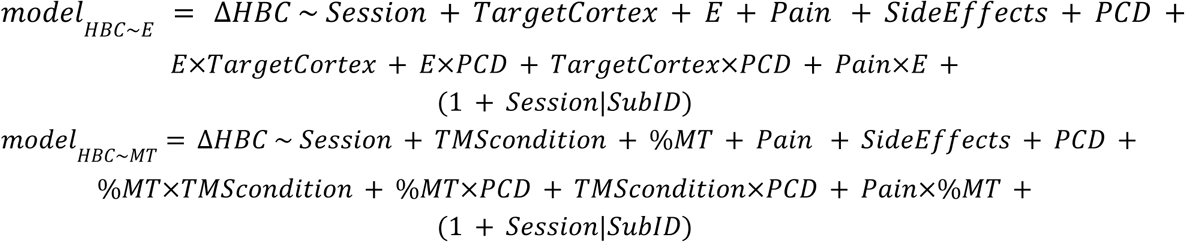

Here, *ΔHBC* denotes the change in HBC relative to stimulation intensity 0%; *TargetCortex* indexes the cortical site at which the E-field was extracted (E-model); *TMScondition* indexes the coil placement (MT-model); %*MT* is stimulator output in percent of motor threshold; *Pain* is the reported pain at the coil placement; *SideEffects* is the summed side-effects score; *PCD* is trial-wise pulsewise coil displacement; and *Session* is the session factor (1–3). All zero-intensity conditions (first cycle) were excluded. In a first step, we restricted E-field values to the targeted cortical site (E-field under the coil: *TargetCortex* = *TMScondition*) to equate the number of observations across models.

Next, we tested whether DLPFC–sgACC FC at the cortical site adds explanatory power by extending both models with FC terms:

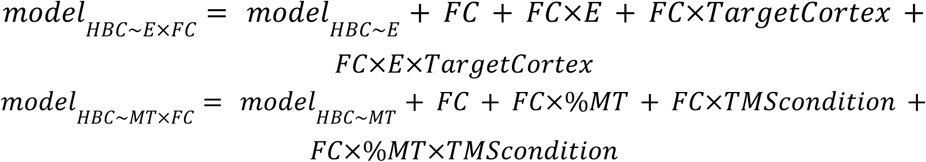

To generalize the E-field perspective beyond “under-coil” tissue, we also fitted the E-only and E-plus-FC models using data from all coil placements for each cortical extraction site (e.g. stimulation at F3 anterior also generates an E-field at F3 lateral and the other sites).

The factors *TMScondition* and *TargetCortex* were encoded using simple-effects contrasts, relative to the 5-cm reference level. In the LMMs, dose, FC and PCD were mean-centred; pain and side effects were standardized. LMM Akaike’s Information Criterion (AIC) comparisons between models with different fixed effects used maximum-likelihood fits on the same rows. Nested dose-only versus dose-plus-FC LMMs were additionally compared using likelihood-ratio tests. Marginal and conditional R² in Table 1 were obtained from REML fits and AIC was chosen as the primary selection metric. Comparisons of the same models fit to different datasets (“E-field under coil” vs “E-field from all coil placements”) relied on R² and inspection of fixed-effect estimates rather than AIC.

**Table 1.** Model comparison to predict ΔHBC.

| Model | AIC | R2<br>(cond.) | R2<br>(marg.) | -2*log(L) | *χ*² | *p* |
|---|---|---|---|---|---|---|
| *MT* | 13048 | 0.168 | 0.061 | 12986 |  |  |
| *MT×FC* | 13005 | 0.188 | 0.077 | 12919 | 66.696 | 1.322 × 10<sup>-9</sup> |
| <sup>*E*</sup> | 13002 | 0.206 | 0.085 | 12940 |  |  |
| *E×FC* | **12948** | **0.236** | **0.104** | **12862** | 78.419 | 8.257 × 10<sup>-12</sup> |

***Notes***: Linear mixed effects models fitted to 3,502 observations from 15 participants. MT: motor threshold; E: cortical E-field; FC: functional connectivity. AIC, likelihoods and FC-module likelihood-ratio tests use ML; marginal and conditional R² use REML. Likelihood-ratio tests have 12 degrees of freedom and nominal, unadjusted P values. Bold indicates the best values.

Model fit indices were obtained with the performance package [47], and significance tests for main effects and factor levels were conducted using lmerTest [48]. Convergence was recorded for all final models. Tabular summaries retained the original sjPlot tab_model layout (v2.9.0) [49] with estimates exported from fitted objects and Satterthwaite degrees of freedom for LMM coefficient tests. Because HBC is known to show nonlinear behavior [12], we additionally fit generalized additive models (GAMs) with analogous predictors with the mgcv package [50]. See SI for full GAM model equation. The matched E-only and full-FC GAM comparisons are provided in Table S1.

As a sensitivity analysis, we refitted the stimulated-target LMMs and GAMs after omitting pain, side effects, PCD and their interactions, using the same observations, core dose/FC terms and participant random effects (Table S2).

For regional controls, we replaced sgACC FC with FC between the same target locations and posterior cingulate cortex (PCC; MNI 0, −53, 26) [51], left/right posterior inferior parietal lobule (IPL; −47, −71, 29 and 50, −64, 27) [52], or left/right occipital seeds of the visual network (MNI −19, −98, 3 and 19, −98, 3) [53]. All seeds were spherical and grey-matter constrained. We retained the same recorded observations, full model formulas and covariates, 10-mm seed/target radii, 3-mm whole-GM FWHM and GSR. Coordinate sources are detailed in the SI (Control-seed analyses). Matched AIC and R² comparisons with bilateral sgACC are reported in Table S3, and single-right sgACC comparisons in Table S4.

#### Voxel-wise mapping of E-field-HBC modulation

As a final generalization step, we mapped the relationship between local E-field strength and HBC on the grey matter surface within the DLPFC ROI for each of the 15 participants. Consistent with previous E-field mapping studies [38, 54, 55], we fitted a linear regression with an intercept at each element, using raw HBC as the response and local E-field strength as the predictor across recorded positive-intensity stimulations. Previous work has shown that motor cortex organization can be mapped reliably with a similar data volume, whereas functions with lower signal-to-noise ratios, such as attention, require more data for stable estimates [55]. Given the high inter- and intra-individual variance in our HBC dataset and the unknown E–HBC input–output function (e.g. linear, sigmoidal, inverted-U), we adopted a simple linear model as a first approximation. For each grey matter ROI element, explained variance was quantified as the squared Pearson correlation between HBC and local E-field strength (R² = r²).

Individual voxel-based sgACC FC maps were projected onto the same DLPFC mid-layer surface. Within each participant, we calculated the Pearson correlation between R² and signed sgACC FC across paired valid DLPFC cortical elements, without applying E-field, R² or FC thresholds. The resulting participant-level correlation coefficients were Fisher z-transformed and tested against zero using a two-sided one-sample t-test. FC maps used the bilateral sgACC seed (10-mm radius), mean seed time series, global signal regression and 8-mm whole-grey-matter FWHM.

## Results

We compared cortical E-field and motor-threshold dose models, examined sgACC FC and regional controls at stimulated targets, and assessed individual surface-map correspondence with sgACC FC.

### Cortical stimulation strength explains heart–brain coupling

We first asked whether HBC modulation is better captured by conventional stimulation intensity in terms of motor threshold or by individual cortical E-field exposure. To this end, we compared two linear mixed-effects models predicting ΔHBC, one using stimulation intensity normalized to each participant’s motor threshold (%MSO, expressed as %MT) and the other using the simulated cortical E-field at the stimulated DLPFC site as the primary dose predictor (Fig. 2). Both models used the same recorded observations, included pain, side effects, and pulsewise coil displacement (PCD) as covariates. The E-field model (*model_HBC_*_∼*E*_) provided a clearly better fit than the MT model (*model_HBC_*_∼*MT*_), with a lower AIC (13,002.15 versus 13,047.76; ΔAIC = AIC_E − AIC_MT = −45.61; Table 1), higher marginal R² (0.085 versus 0.061) and higher conditional R² (0.206 versus 0.168). Under conventional AIC interpretation [56, 57], differences greater than 10 indicate essentially no support for the inferior model; the observed AIC difference of 45.6 corresponds to an evidence ratio of approximately 8 × 10⁹ in favour of the E-field model, indicating that stimulator-output-based dosing is negligible as a competing description of the same data. Thus, for the same experimental data, individual cortical E-field estimates captured more variance in ΔHBC than motor-threshold-based intensity, supporting cortical E-field exposure as a more informative dose descriptor for HBC modulation.

**Figure 2.**
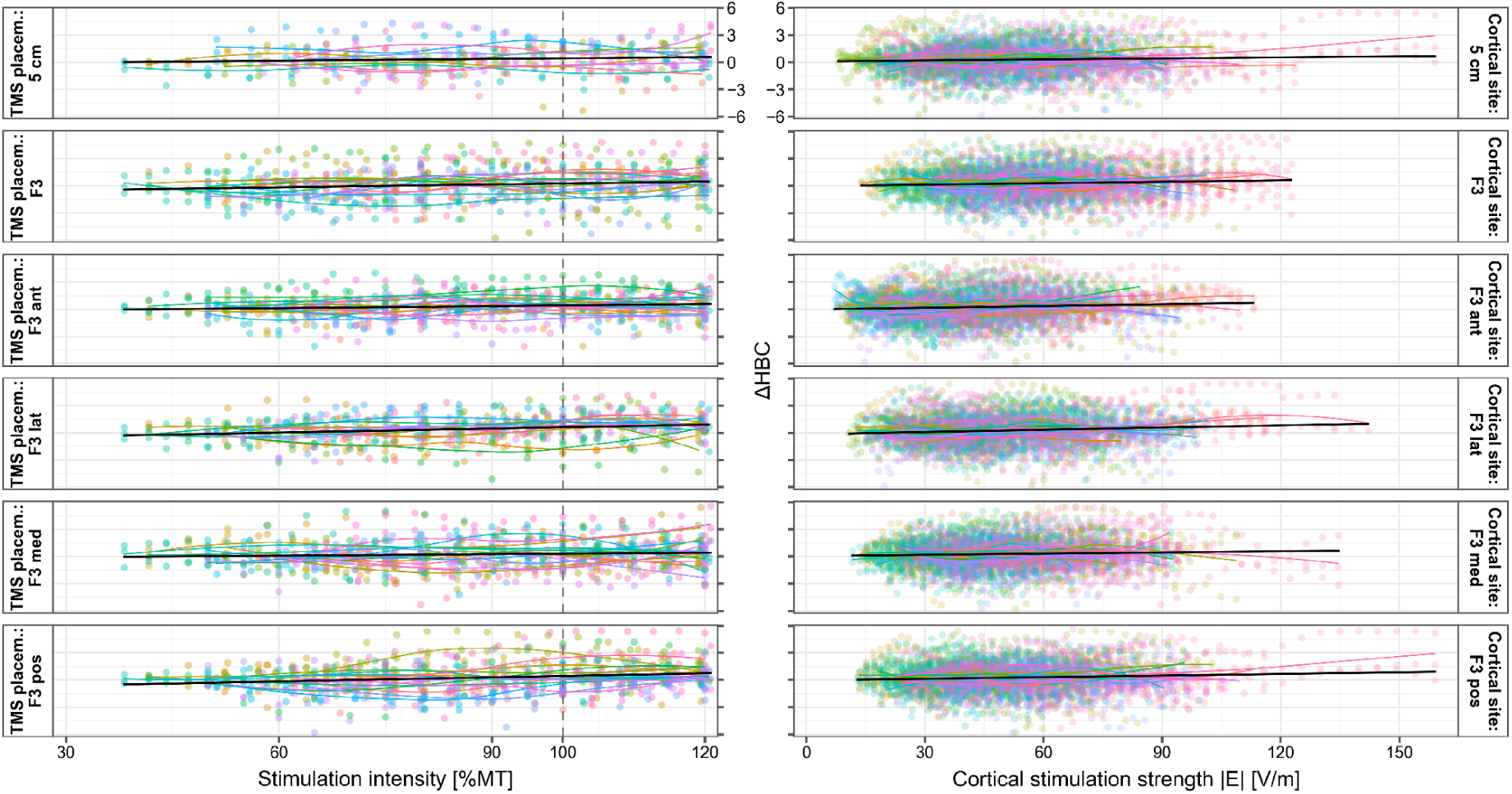
Heart–brain coupling in relation to stimulator output and cortical E-field exposure. Left: In the conventional coil-centered view, stimulation targets are defined by TMS coil placements (rows) and stimulation intensity relative to individual motor threshold (%MT). For each TMS burst per intensity and coil placement, one ΔHBC data point is obtained. Right: In the E-field based analyses stimulation targets are defined as cortical sites (rows), and dose is quantified as cortical E-field magnitude (V/m). Because placing the TMS coil above one cortical site also yields cortical stimulation of the other targets, this allows integrating information from multiple coil placements. The same experimental data are shown in both columns. Colors indicate individual subjects. Black lines show pooled linear fits and coloured curves show participant-specific LOESS fits. ΔHBC: HBC change defined as log(HBC_intX_) - log(HBC_int0_).

### Functional connectivity modulates heart–brain coupling

We next asked whether individual differences in DLPFC–sgACC connectivity further explain variance in ΔHBC beyond cortical E-field and nuisance covariates. Adding FC and its interactions to the stimulated-target E-field LMM improved model fit, reducing AIC by 54.42 (likelihood-ratio χ²(12) = 78.42, *p* = 8.26 × 10⁻¹²). Marginal R² increased from 0.085 to 0.104 and conditional R² from 0.206 to 0.236 (Table 1; Tables S5–S7). In the corresponding GAM, adding the full FC module reduced AIC by 499.21 and increased adjusted R² from 0.194 to 0.316 (Table S1). Thus, sgACC connectivity provided information about HBC modulation beyond that captured by cortical stimulation strength alone.

The E-field advantage and FC contribution persisted after omitting pain, side effects, PCD and their interactions. The E-field LMM retained a 43.14-unit AIC advantage over MT, and adding FC improved both the LMM (χ²(12) = 88.26, p = 1.07 × 10⁻¹³) and GAM (AIC reduction = 594.93, Table S2).

We then examined how the E-field, FC, and cortical site jointly shape HBC. In the stimulated-target E-plus-FC LMM, the FC×E×TargetCortex contrasts relative to the 5-cm reference were negative at F3 and F3 lateral (*p* < 0.05; Table S8). These interactions indicate site differences in how sgACC connectivity modifies the E-field–HBC relationship. Sensitivity analyses across extraction radii and FC smoothing kernels are reported in Tables S9–S12. PCD-related effects also varied across cortical sites, with significant PCD-by-target contrasts in the LMM and site-specific PCD smooths in the GAM (Table S13).

Within the full models, we visualized the predicted E-field–ΔHBC relationships at negative FC (−0.1) and positive FC (+0.1) across stimulated targets (Fig. 3). At F3 and F3 lateral, both the LMM and GAM predicted greater ΔHBC at higher E-field strengths for negative than for positive FC. The GAM additionally revealed site-dependent nonlinear profiles. At F3, the positive-FC curve reached a local predicted maximum at approximately 65–70 V/m and declined at higher field strengths, whereas the negative-FC curve increased over the higher-dose range. Negative-FC predictions also increased across the displayed E-field range at F3 anterior and F3 lateral. These profiles illustrate how cortical site and sgACC connectivity jointly shape the predicted stimulation–response relationship. Figure S1 summarizes FC variation across smoothing kernels; the six-location model predictions are presented descriptively in Fig. S2. Importantly, besides showing specificity of the different DLPFC targets with sgACC, we also tested for potential effects of DLPFC connectivity to other brain targets: PCC, left and right IPL, and left and right occipital seeds of the visual network. Specifically, we reused the same linear and nonlinear model equations as before with FC values to these targets. Among all tested regions, sgACC yielded the lowest stimulated-target GAM AIC (12,313.35; adjusted R² = 0.316); the closest control was the right occipital visual seed, with a ΔAIC of +13.91. The sgACC model also had the lowest stimulated-target LMM AIC (12,947.73), followed by right occipital visual FC (12,958.42; ΔAIC = 10.69; Table S3). Corresponding single-right-seed results are reported in Table S4.

**Figure 3.**
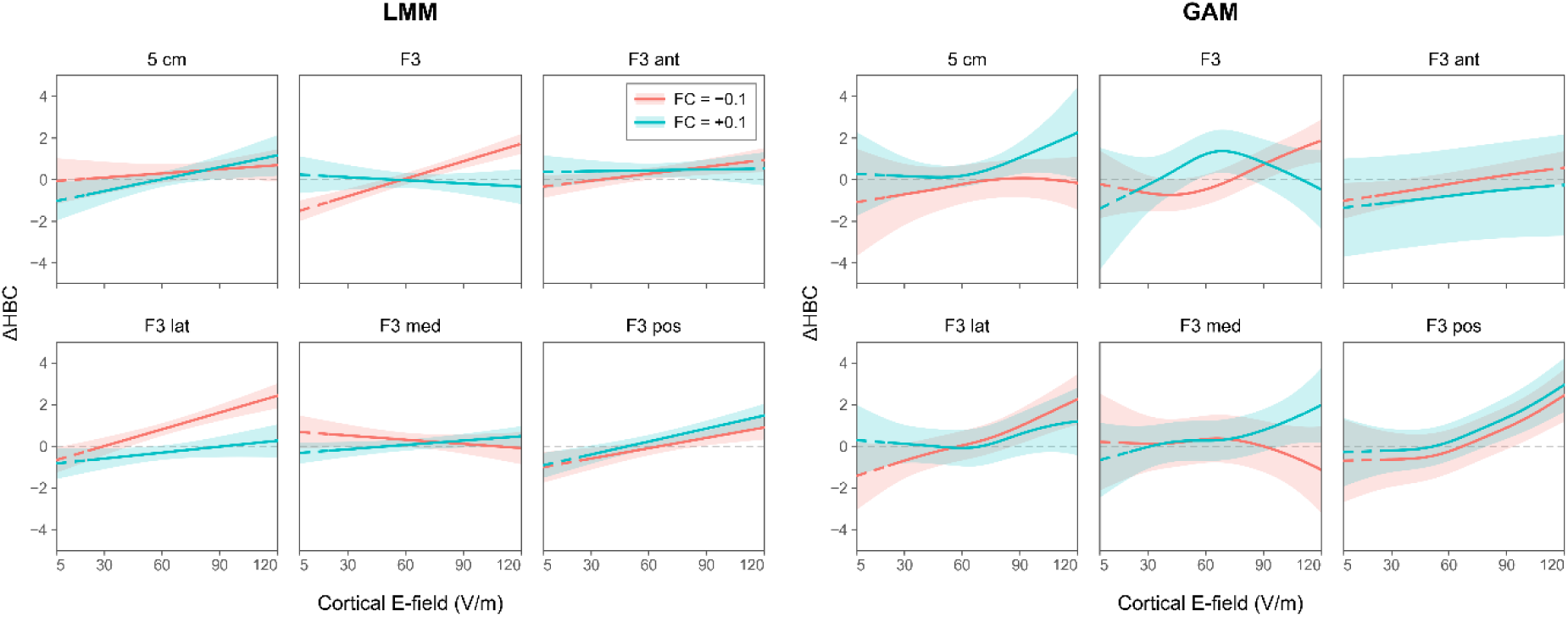
Linear and nonlinear models link cortical stimulation to heart–brain coupling. Stimulated-target predictions from the full LMM (left; REML) and GAM (right; fREML) fitted to 3,502 responses from 15 participants are shown for each cortical site. Both models use the bilateral sgACC seed, mean seed time series, median target FC, 10-mm seed/target radii, 3-mm whole-grey-matter FWHM and GSR. Curves show FC = −0.1 (red) and +0.1 (cyan) over E = 5–120 V/m. Shaded bands indicate pointwise 95% confidence intervals. Dashed segments mark E-field or FC values outside the site-specific observed marginal ranges. ΔHBC = log(HBC_intX_) − log(HBC_int0_).

### Voxel-wise mapping of HBC modulation

After establishing site-level effects of cortical stimulation strength and sgACC connectivity, we investigated whether these relationships extended to a finer spatial scale within the DLPFC. By combining E-field estimates across coil placements, we sought to identify cortical locations where variation in local stimulation strength most closely tracked HBC. For each participant, we therefore regressed raw HBC on the local E-field at each element of the DLPFC surface ROI across recorded coil placements and positive stimulation intensities (Fig. 4). Extending previous E-field mapping approaches in motor and parietal cortex [38, 54, 55], this analysis generated individual maps of the proportion of HBC variance explained by local cortical E-field exposure.

**Figure 4.**
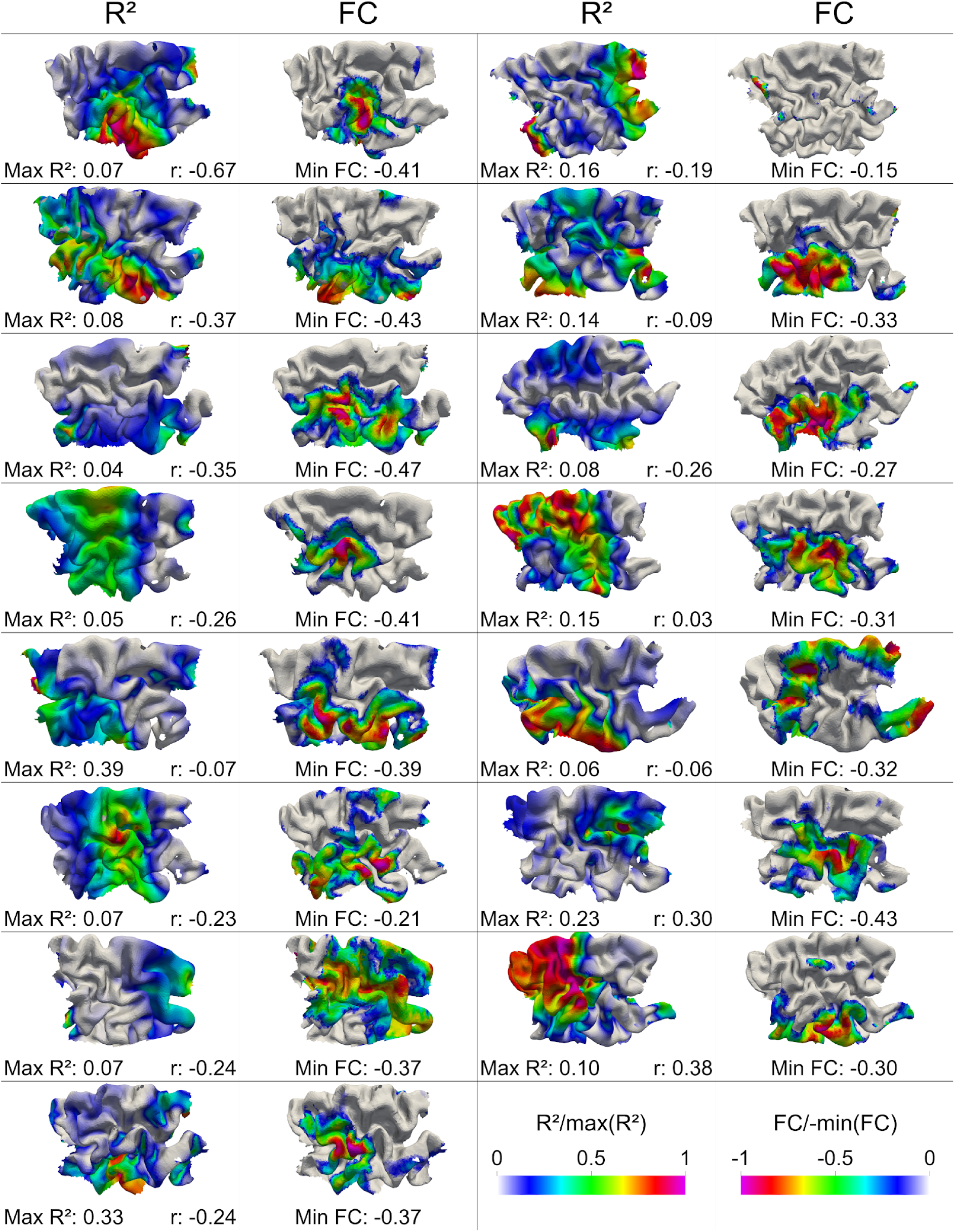
Spatial maps of HBC explained variance and sgACC functional connectivity. For each element of the DLPFC surface ROI, raw heart–brain coupling (HBC) was regressed on local E-field across recorded positive-intensity stimulations, separately for each participant (columns “R²”). The coefficient of determination quantifies the fraction of HBC variance explained at each cortical element. Voxel-based FC maps use the bilateral sgACC seed (10-mm radius), mean seed time series, GSR and 8-mm whole-grey-matter FWHM. For display, R² is divided by the participant-specific maximum R², and negative FC is divided by the absolute participant-specific minimum FC (columns “FC”). The two colour scales use the same palette in opposite directions: warmer colours indicate higher normalized R² and more negative normalized FC. Spatial correlations *r*eported in the text use paired valid elements and all signed, unnormalized FC values, without E-field, R² or FC thresholds.

Explained variance varied across cortical locations and participants, with participant-specific maxima differing across the 15 individuals. Regions with higher R² overlapped spatially with DLPFC areas showing negative sgACC connectivity. To quantify this correspondence, we calculated each participant’s spatial correlation between unthresholded R² and signed sgACC FC across paired valid cortical elements. The Fisher-z-transformed correlations were significantly negative at the group level (*t*(14) = −2.242, *p* = 0.0417). This spatial correspondence complements the site-level models by linking the cortical distribution of E-field–HBC associations to individual DLPFC–sgACC connectivity.

Complementing these explained-variance maps, element-wise dose–response gain maps (absolute regression slopes of HBC on local E-field; Fig. S3) closely matched the R² maps: after excluding elements with E-field variance below the participant-specific 25th percentile, R² and absolute gain correlated at r = 0.84 ± 0.03 (mean ± SD) across participants (similar for other percentiles). Thus, regions of high E-field–HBC predictability coincided with regions of strong stimulation–response coupling.

## Discussion

TMS of the DLPFC is an established treatment for depression, yet its mechanistic effects remain unclear. A prominent hypothesis is the engagement of frontal–vagal circuitry via DLPFC–sgACC connectivity, thus shaping autonomic regulation. Here, we provide strong support for this hypothesis. Modeling the acute neuro-cardiac response to DLPFC stimulation within an individualized, cortex-centered framework, we found that both cortical dose and network embedding matter: cortical E-field strength explained more HBC variance than motor-threshold-based intensity, sgACC connectivity further improved model fit with steeper predicted responses at negatively connected targets, and individual maps of E-field–HBC coupling spatially overlapped with sgACC-anticorrelated cortex. Together, these results show that the cardiac response to prefrontal TMS is determined by the cortical exposure actually delivered and the autonomic network connected to it. These findings support HBC as a network-sensitive physiological readout and provide a basis for cortical and network-informed dosing studies.

### E-field-based dosing and network embedding

Our site-level analyses demonstrated that defining TMS dose at the cortical level captured more HBC variance than motor-threshold-based intensity, despite identical random-effects structures and covariates. This pattern suggests that some variability traditionally attributed to noise or inter-individual differences in excitability may relate to differences in cortical stimulation exposure, consistent with previous work on E-field-based dosing [24, 25]. Adding DLPFC–sgACC connectivity and its interactions with E-field and cortical site further improved model performance, indicating that predicted HBC responses at a given E-field depend on the network context of the stimulated tissue.

At the stimulated targets, both linear and nonlinear models favoured FC with sgACC over the control regions PCC, IPL, and occipital visual seeds. The site-specific responses converge with connectivity-guided antidepressant targeting [13, 14, 17], the value of defining stimulated tissue using E-fields [18], and evidence for distinct directed connectivity patterns among prefrontal subregions [20, 21]. More broadly, the connectivity-moderated dose–response relationships we identified offer a possible explanation for why the literature has not converged on a single optimal stimulation intensity [58, 59]: the same nominal dose engages different cortical exposure across individuals [23, 60], and the resulting input–output functions are non-monotonic and site-dependent [61–63], such that studies differ in which segment of a moderated dose–response curve they sample. Our findings extend this framework to physiological responses during prefrontal stimulation.

### From focal targets to distributed maps

Our high-resolution cortical mapping extends the stimulation-site-level findings to spatially resolved, individual cortical maps. By regressing HBC on the local E-field at each DLPFC cortical element, we identified locations where variation in cortical stimulation strength most closely tracked HBC, extending previous high-resolution TMS mapping approaches in motor and parietal cortex [38, 54, 55]. Regions of higher explained variance varied across individuals and overlapped with negative sgACC connectivity, adding a spatial dimension to the site-level evidence: it is not only the predefined targets, but the surrounding cortical neighborhood, where E-field–HBC coupling aligns with sgACC-anticorrelated tissue. This spatial correspondence is consistent with the DLPFC-sgACC-vagus nerve model of autonomic regulation and complements the site-level evidence by linking HBC modulation to sgACC connectivity within individual anatomy rather than group-defined coordinates.

### Translational implications for neuro-cardiac TMS

These associations motivate clinical studies of individualized cortical dosing for autonomic regulation. Such studies could compare E-field-calibrated intensity with conventional %MT dosing and examine whether connectivity-dependent HBC profiles help explain heterogeneous treatment responses (e.g. Dijkstra et al. [64]). Evidence that DLPFC stimulation evokes sgACC responses [65, 66] further motivates testing HBC as an accessible physiological readout of frontal–sgACC engagement. Where individual FC mapping is unavailable, population-based connectivity targets and online HBC could be evaluated for guiding dose adjustment [19].

Combining individual E-fields, connectivity and physiological readouts also offers a spatial mapping strategy for studying association cortex. Applied prospectively, this approach may help evaluate where and how strongly to stimulate connectivity-defined DLPFC regions to influence autonomic and depressive outcomes [18].

### Methodological considerations and limitations

Coil displacement remains relevant when relating static E-field simulations to changing stimulation conditions. Including pulsewise coil displacement (PCD, Numssen et al. [43]) and other targeting-precision metrics [67, 68] helps characterize the relationship between simulated cortical dose and HBC under variable coil positioning. FC estimation also remains subject to resting-state reliability constraints and susceptibility-related signal loss in ventral prefrontal regions [69, 70].

Several further limitations merit consideration. The present study used a relatively small sample of healthy adults and focused on acute HBC responses rather than clinical or long-term autonomic outcomes, limiting direct generalizability to MDD treatment. HBC captures frequency-locked modulation at the TMS cycle, and other aspects of autonomic regulation (e.g. slower HRV components) may show different spatial and dosing profiles. The spatial mapping analyses relied on linear models because the true E-field–HBC input–output function is unknown and our ability to detect more complex shapes is constrained by signal-to-noise and data density. Finally, while the spatial mapping analyses reveal distributed patterns within DLPFC, they do not capture potential downstream subcortical or brainstem contributions that are not directly modelled in the cortical E-field.

## Conclusions

Together with prior Neuro-Cardiac-Guided TMS [11, 19, 64] and trigeminal-control studies [71], our results support HBC as a candidate physiological marker for studying frontal–vagal function. We extend this framework by showing that HBC responses vary with both local cortical E-fields and sgACC-centred connectivity, situating HBC within a cortex- and network-based approach to prefrontal stimulation.

Future work should extend this connectivity-informed, E-field-based framework to clinical populations [64] and longitudinal treatment protocols. A key question is whether sgACC-anticorrelated DLPFC regions showing higher E-field–HBC explained variance also yield greater symptomatic improvement when stimulation is dosed using individual E-fields rather than %MT. Integrating online HBC with individual FC maps, while accounting for state-dependent stimulation effects [72], could support adaptive protocols that adjust target, intensity, and timing. Applying the same mapping approach to other targets and readouts, including cognitive control, affective processing and sleep, may further clarify how cortical stimulation engages networks underlying physiological and behavioural responses.

## Data availability

Data and analysis materials for the present study are available via an OSF project: https://osf.io/v9c2t. This repository contains HBC data, E-field data in the DLPFC ROI, DLPFC–sgACC FC, side effects and scripts to reproduce the analyses and figures reported here. Analysis code is available at https://gitlab.gwdg.de/tms-localization/papers/tms-hbc-efield-connectivity. ECG preprocessing outputs and HBC measures from the previous NCG-TMS study are available at https://gitlab.gwdg.de/tms-localization/papers/tms-hbc.

## Acknowledgement

This work was supported by Lise Meitner Excellence funding from the Max Planck Society, the European Research Council (ERC-2021-COG 101043747), and the German Research Foundation (HA 6314/3-1, HA 6314/4-2, HA 6314/9-1, Research Unit 5429/1 (467143400: HA 6314/10-1). Ole Numssen and Konstantin Weise were supported by the Federal Ministry of Research, Technology and Space (Bundesministerium für Forschung, Technologie und Raumfahrt, BMFTR, Grant no. 01GQ2201 to Thomas Knösche). Ole Numssen was supported by ScaDS.AI.

## Author contributions

Z.F., S.M., K.W., G.H., T.K. and O.N. conceived the study. Z.F., S.M. and O.N. developed the methodology. Z.F. and G.G. collected and curated the data. Z.F., S.M., G.G. and O.N. performed the formal analyses. Z.F., S.M., K.W. and O.N. developed the analysis software. Z.F. and O.N. verified the analyses and results. Z.F., S.M. and O.N. prepared the figures and wrote the initial manuscript. Z.F., S.M., K.W., G.H., T.K. and O.N. critically reviewed and revised the manuscript. G.H., T.K. and O.N. supervised the research. G.H. and T.K. acquired funding, provided resources and coordinated the project.

## References

1. Berger S, Kliem A, Yeragani V, Bar KJ. Cardio-respiratory coupling in untreated patients with major depression. J Affect Disord 2012; 139: 166–171.

2. Thayer JF, Ahs F, Fredrikson M, Sollers JJ, 3rd, Wager TD. A meta-analysis of heart rate variability and neuroimaging studies: implications for heart rate variability as a marker of stress and health. Neurosci Biobehav Rev 2012; 36: 747–756.

3. Thayer JF, Lane RD. Claude Bernard and the heart-brain connection: further elaboration of a model of neurovisceral integration. Neurosci Biobehav Rev 2009; 33: 81–88.

4. Makovac E, Thayer JF, Ottaviani C. A meta-analysis of non-invasive brain stimulation and autonomic functioning: Implications for brain-heart pathways to cardiovascular disease. Neurosci Biobehav Rev 2017; 74: 330–341.

5. Schmausser M, Hoffmann S, Raab M, Laborde S. The effects of noninvasive brain stimulation on heart rate and heart rate variability: A systematic review and meta-analysis. J Neurosci Res 2022; 100: 1664–1694.

6. Morriss R, Webster L, Ingram L, Abdelghani M, Anton A, Barber S et al. Connectivity guided intermittent theta burst stimulation versus repetitive transcranial magnetic stimulation in moderately severe treatment resistant depression. Efficacy and Mechanism Evaluation. Southampton (UK) 2025.

7. Rajasekharan D, Madore MR, Holtzheimer P, Lim KO, Williams LM, Philip NS. Personalized models of Beam/F3 targeting in transcranial magnetic stimulation for depression: Implications for precision clinical translation. Brain Stimul 2025; 18: 829–837.

8. Iseger TA, Padberg F, Kenemans JL, Gevirtz R, Arns M. Neuro-Cardiac-Guided TMS (NCG-TMS): Probing DLPFC-sgACC-vagus nerve connectivity using heart rate - First results. Brain Stimul 2017; 10: 1006–1008.

9. Iseger TA, Padberg F, Kenemans JL, van Dijk H, Arns M. Neuro-Cardiac-Guided TMS (NCG TMS): A replication and extension study. Biol Psychol 2021; 162: 108097.

10. Iseger TA, van Bueren NER, Kenemans JL, Gevirtz R, Arns M. A frontal-vagal network theory for Major Depressive Disorder: Implications for optimizing neuromodulation techniques. Brain Stimul 2020; 13: 1–9.

11. Dijkstra E, van Dijk H, Vila-Rodriguez F, Zwienenberg L, Rouwhorst R, Coetzee JP et al. Transcranial Magnetic Stimulation-Induced Heart-Brain Coupling: Implications for Site Selection and Frontal Thresholding-Preliminary Findings. Biol Psychiatry Glob Open Sci 2023; 3: 939–947.

12. Feng ZJ, Martin S, Numssen O, Weise K, Jing Y, Gerardos G et al. Target-Specificity and repeatability in neuro-cardiac-guided TMS for heart-brain coupling. Transl Psychiatry 2026; 16:

13. Fox MD, Buckner RL, White MP, Greicius MD, Pascual-Leone A. Efficacy of transcranial magnetic stimulation targets for depression is related to intrinsic functional connectivity with the subgenual cingulate. Biol Psychiatry 2012; 72: 595–603.

14. Weigand A, Horn A, Caballero R, Cooke D, Stern AP, Taylor SF et al. Prospective Validation That Subgenual Connectivity Predicts Antidepressant Efficacy of Transcranial Magnetic Stimulation Sites. Biol Psychiat 2018; 84: 28–37.

15. Cash RFH, Weigand A, Zalesky A, Siddiqi SH, Downar J, Fitzgerald PB et al. Using Brain Imaging to Improve Spatial Targeting of Transcranial Magnetic Stimulation for Depression. Biol Psychiatry 2021; 90: 689–700.

16. Cash RFH, Zalesky A, Thomson RH, Tian Y, Cocchi L, Fitzgerald PB. Subgenual Functional Connectivity Predicts Antidepressant Treatment Response to Transcranial Magnetic Stimulation: Independent Validation and Evaluation of Personalization. Biol Psychiatry 2019; 86: e5–e7.

17. Cole EJ, Phillips AL, Bentzley BS, Stimpson KH, Nejad R, Barmak F et al. Stanford Neuromodulation Therapy (SNT): A Double-Blind Randomized Controlled Trial. Am J Psychiatry 2022; 179: 132–141.

18. Elbau IG, Lynch CJ, Downar J, Vila-Rodriguez F, Power JD, Solomonov N et al. Functional Connectivity Mapping for rTMS Target Selection in Depression. Am J Psychiatry 2023; 180: 230–240.

19. Dijkstra ESA, Frandsen SB, van Dijk H, Duecker F, Taylor JJ, Sack AT et al. Probing prefrontal-sgACC connectivity using TMS-induced heart–brain coupling. Nature Mental Health 2024; 2: 809–817.

20. Avalos-Alais S, Jedynak M, Boyer A, Chanteloup-Foret B, Pinheiro C, Cline CC et al. High-resolution electrophysiological mapping of effective connectivity of lateral prefrontal cortex. Brain 2026; 149: 963–975.

21. Batail JM, Borgmann L. Mapping the human lateral prefrontal cortex at the circuit level. Brain 2026; 149: 707–709.

22. Siddiqi SH, Philip NS. Hitting the Target of Image-Guided Psychiatry? Am J Psychiatry 2023; 180: 185–187.

23. Soleimani G, Alekseichuk I, Aurup C, Bergmann TO, Bestmann S, Beynel L et al. Dose-response relationships in transcranial brain stimulation: Physics, physiology and mechanism. Brain Stimul 2026; 19: 103067.

24. Dannhauer M, Gomez LJ, Robins PL, Wang D, Hasan NI, Thielscher A et al. Electric Field Modeling in Personalizing Transcranial Magnetic Stimulation Interventions. Biol Psychiatry 2024; 95: 494–501.

25. Numssen O, Kuhnke P, Weise K, Hartwigsen G. Electric-field-based dosing for TMS. Imaging Neurosci (Camb) 2024; 2: 1–12.

26. Stokes MG, Chambers CD, Gould IC, English T, McNaught E, McDonald O et al. Distance-adjusted motor threshold for transcranial magnetic stimulation. Clin Neurophysiol 2007; 118: 1617–1625.

27. Thielscher A, Opitz A, Windhoff M. Impact of the gyral geometry on the electric field induced by transcranial magnetic stimulation. Neuroimage 2011; 54: 234–243.

28. Weise K, Numssen O, Kalloch B, Zier AL, Thielscher A, Haueisen J et al. Precise motor mapping with transcranial magnetic stimulation. Nat Protoc 2023; 18: 293–318.

29. Caulfield KA, Li X, George MS. Four electric field modeling methods of Dosing Prefrontal Transcranial Magnetic Stimulation (TMS): Introducing APEX MT dosimetry. Brain Stimul 2021; 14: 1032–1034.

30. Oldfield RC. The assessment and analysis of handedness: the Edinburgh inventory. Neuropsychologia 1971; 9: 97–113.

31. Rossi S, Antal A, Bestmann S, Bikson M, Brewer C, Brockmoller J et al. Safety and recommendations for TMS use in healthy subjects and patient populations, with updates on training, ethical and regulatory issues: Expert Guidelines. Clin Neurophysiol 2021; 132: 269–306.

32. Rossini PM, Burke D, Chen R, Cohen LG, Daskalakis Z, Di Iorio R et al. Non-invasive electrical and magnetic stimulation of the brain, spinal cord, roots and peripheral nerves: Basic principles and procedures for routine clinical and research application. An updated report from an I.F.C.N. Committee. Clin Neurophysiol 2015; 126: 1071–1107.

33. Giustiniani A, Vallesi A, Oliveri M, Tarantino V, Ambrosini E, Bortoletto M et al. A questionnaire to collect unintended effects of transcranial magnetic stimulation: A consensus based approach. Clin Neurophysiol 2022; 141: 101–108.

34. Esteban O, Markiewicz CJ, Blair RW, Moodie CA, Isik AI, Erramuzpe A et al. fMRIPrep: a robust preprocessing pipeline for functional MRI. Nat Methods 2019; 16: 111–116.

35. Abraham A, Pedregosa F, Eickenberg M, Gervais P, Mueller A, Kossaifi J et al. Machine learning for neuroimaging with scikit-learn. Front Neuroinform 2014; 8: 14.

36. Sun W, Billot A, Du J, Wei X, Lemley RA, Daneshzand M et al. Precision Network Modeling of Transcranial Magnetic Stimulation Across Individuals Suggests Therapeutic Targets and Potential for Improvement. medRxiv 2024;

37. Wang HT, Meisler SL, Sharmarke H, Clarke N, Gensollen N, Markiewicz CJ et al. Continuous evaluation of denoising strategies in resting-state fMRI connectivity using fMRIPrep and Nilearn. PLoS Comput Biol 2024; 20: e1011942.

38. Numssen O, Zier AL, Thielscher A, Hartwigsen G, Knosche TR, Weise K. Efficient high-resolution TMS mapping of the human motor cortex by nonlinear regression. Neuroimage 2021; 245: 118654.

39. Fox MD, Liu H, Pascual-Leone A. Identification of reproducible individualized targets for treatment of depression with TMS based on intrinsic connectivity. Neuroimage 2013; 66: 151–160.

40. Puonti O, Van Leemput K, Saturnino GB, Siebner HR, Madsen KH, Thielscher A. Accurate and robust whole-head segmentation from magnetic resonance images for individualized head modeling. Neuroimage 2020; 219: 117044.

41. Thielscher A, Antunes A, Saturnino GB. Field modeling for transcranial magnetic stimulation: A useful tool to understand the physiological effects of TMS? Annu Int Conf IEEE Eng Med Biol Soc 2015; 2015: 222–225.

42. Fischl B. FreeSurfer. Neuroimage 2012; 62: 774–781.

43. Numssen O, Martin S, Williams K, Knosche TR, Hartwigsen G. Quantification of subject motion during TMS via pulsewise coil displacement. Brain Stimul 2024; 17: 1045–1047.

44. Van Hoornweder S, Geraerts M, Verstraelen S, Nuyts M, Caulfield KA, Meesen R. Differences in scalp-to-cortex tissues across age groups, sexes and brain regions: Implications for neuroimaging and brain stimulation techniques. Neurobiol Aging 2024; 138: 45–62.

45. Van Hoornweder S, Nuyts M, Frieske J, Verstraelen S, Meesen RLJ, Caulfield KA. Outcome measures for electric field modeling in tES and TMS: A systematic review and large-scale modeling study. Neuroimage 2023; 281: 120379.

46. Bates D, Mächler M, Bolker B, Walker S. Fitting Linear Mixed-Effects Models Usinglme4. Journal of Statistical Software 2015; 67: 1–48.

47. Lüdecke D, Ben-Shachar M, Patil I, Waggoner P, Makowski D. performance: An R Package for Assessment, Comparison and Testing of Statistical Models. Journal of Open Source Software 2021; 6:

48. Kuznetsova A, Brockhoff PB, Christensen RH. lmerTest package: tests in linear mixed effects models. Journal of statistical software 2017; 82: 1–26.

49. Lüdecke D. sjPlot-Data Visualization for Statistics in Social Science. Zenodo 2021;

50. Wood SN. Generalized additive models: an introduction with R. chapman and hall/CRC; 2017.

51. Van Dijk KR, Hedden T, Venkataraman A, Evans KC, Lazar SW, Buckner RL. Intrinsic functional connectivity as a tool for human connectomics: theory, properties, and optimization. J Neurophysiol 2010; 103: 297–321.

52. Vincent JL, Kahn I, Snyder AZ, Raichle ME, Buckner RL. Evidence for a frontoparietal control system revealed by intrinsic functional connectivity. J Neurophysiol 2008; 100: 3328–3342.

53. Philippi CL, Pujara MS, Motzkin JC, Newman J, Kiehl KA, Koenigs M. Altered resting-state functional connectivity in cortical networks in psychopathy. J Neurosci 2015; 35: 6068–6078.

54. Vetter DE, Jooss A, Mutanen TP, Kozak G, Ziemann U. Short-interval interhemispheric inhibition does not originate from the motor hotspot. Brain Stimul 2025; 18: 1074–1081.

55. Jing Y, Numssen O, Weise K, Kalloch B, Buchberger L, Haueisen J et al. Modeling the effects of transcranial magnetic stimulation on spatial attention. Phys Med Biol 2023; 68:

56. Burnham KP, Anderson DR, Huyvaert KP. AIC model selection and multimodel inference in behavioral ecology: some background, observations, and comparisons. Behavioral ecology and sociobiology 2011; 65: 23–35.

57. Burnham KP, Anderson DR. Multimodel inference: understanding AIC and BIC in model selection. Sociological methods & research 2004; 33: 261–304.

58. Hsu TW, Yeh TC, Kao YC, Thompson T, Brunoni AR, Carvalho AF et al. The dose-effect relationship of six stimulation parameters with rTMS over left DLPFC on treatment-resistant depression: A systematic review and meta-analysis. Neurosci Biobehav Rev 2024; 162: 105704.

59. Dalhuisen I, van Bronswijk S, Bors J, Smit F, Spijker J, Tendolkar I et al. The association between sample and treatment characteristics and the efficacy of repetitive transcranial magnetic stimulation in depression: A meta-analysis and meta-regression of sham-controlled trials. Neurosci Biobehav Rev 2022; 141: 104848.

60. Caulfield KA, Li X, George MS. A reexamination of motor and prefrontal TMS in tobacco use disorder: Time for personalized dosing based on electric field modeling? Clin Neurophysiol 2021; 132: 2199–2207.

61. Lee JC, Corlier J, Wilson AC, Tadayonnejad R, Marder KG, Ngo D et al. Subthreshold stimulation intensity is associated with greater clinical efficacy of intermittent theta-burst stimulation priming for Major Depressive Disorder. Brain Stimul 2021; 14: 1015–1021.

62. Chung SW, Rogasch NC, Hoy KE, Sullivan CM, Cash RFH, Fitzgerald PB. Impact of different intensities of intermittent theta burst stimulation on the cortical properties during TMS-EEG and working memory performance. Hum Brain Mapp 2018; 39: 783–802.

63. Quinn DK, Upston J, Jones TR, Gibson BC, Olmstead TA, Yang J et al. Electric field distribution predicts efficacy of accelerated intermittent theta burst stimulation for late-life depression. Front Psychiatry 2023; 14: 1215093.

64. Dijkstra ESA, Rouwhorst R, Zwienenberg L, van Oostrom I, van Dijk H, Sack AT et al. TMS-induced heart-brain coupling associated with early clinical response in depression. Brain Stimul 2025; 18: 1744–1746.

65. Solomon EA, Hassan U, Trapp NT, Boes AD, Keller CJ. DLPFC TMS suppresses high-frequency neural activity in the human sgACC. Brain Stimul 2026; 19: 102985.

66. Trapp NT, Liu X, Li Z, Bruss J, Keller CJ, Boes AD et al. Dorsolateral prefrontal cortex TMS evokes responses in the subgenual anterior cingulate cortex: Intracranial EEG evidence from two human cases. Brain Stimul 2025; 18: 1202–1204.

67. Burns MR, Hermiller MS. Quantifying and reporting the precision of transcranial magnetic stimulation targeting. Brain Res 2025; 1849: 149350.

68. Grosshagauer S, Woletz M, Becher M, Bjorklund J, Padberg F, Keeser D et al. Reducing target E-field variability in repetitive TMS through online motion compensation. Brain Stimul 2026; 19: 102990.

69. Devlin JT, Russell RP, Davis MH, Price CJ, Wilson J, Moss HE et al. Susceptibility-induced loss of signal: comparing PET and fMRI on a semantic task. Neuroimage 2000; 11: 589–600.

70. Weiskopf N, Hutton C, Josephs O, Turner R, Deichmann R. Optimized EPI for fMRI studies of the orbitofrontal cortex: compensation of susceptibility-induced gradients in the readout direction. MAGMA 2007; 20: 39–49.

71. Arns M, Williams NR, Downar J, Dojnov A, Coetzee J, Goya-Maldonado R. Trigeminal nerve stimulation as an active control condition in TMS clinical trials: Evidence from heart-brain coupling and clinical outcomes. Brain Stimul 2025; 18: 1403–1406.

72. Grosshagauer S, Woletz M, Vasileiadi M, Linhardt D, Nohava L, Schuler AL et al. Chronometric TMS-fMRI of personalized left dorsolateral prefrontal target reveals state-dependency of subgenual anterior cingulate cortex effects. Mol Psychiatry 2024; 29: 2678–2688.

