## Supplementary material for "Cortical Stimulation Strength and sgACC Connectivity Shape Neuro-Cardiac Responses to Prefrontal TMS": SI

#### Functional connectivity

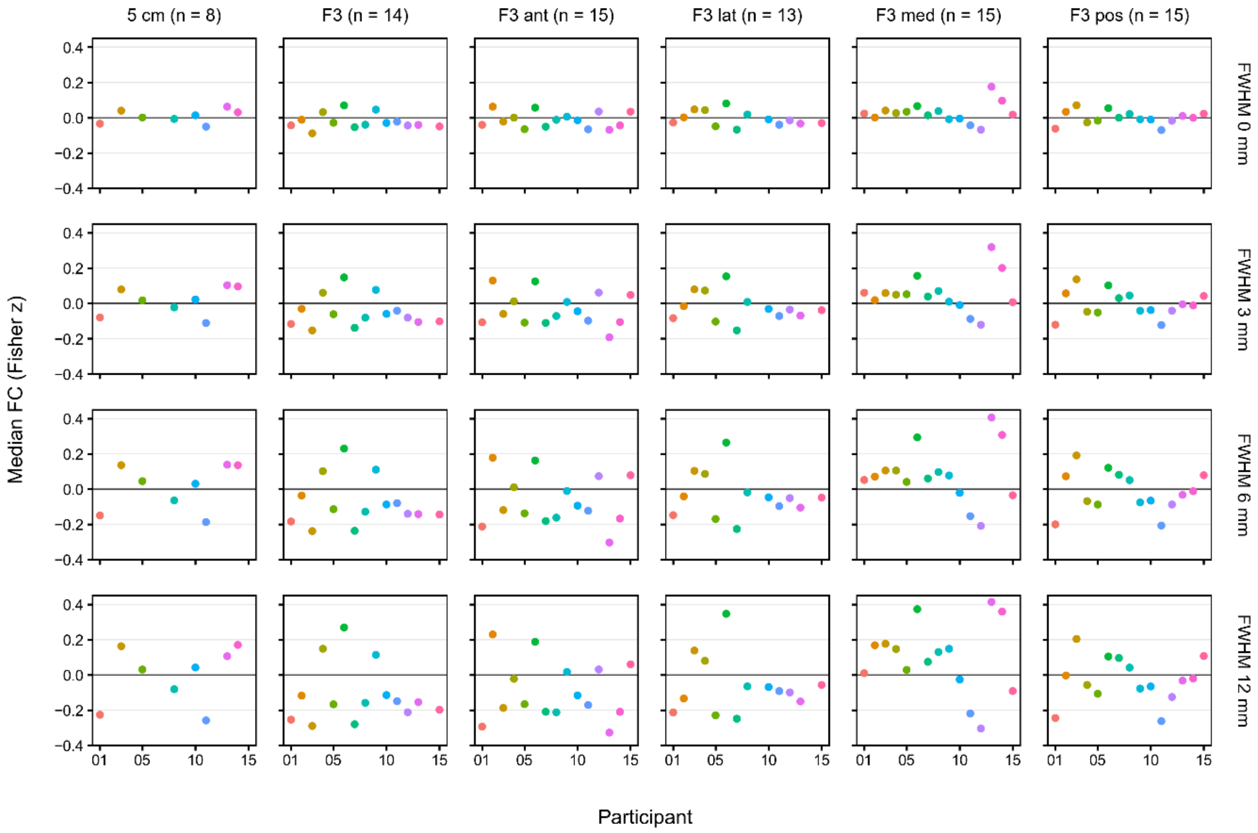


**Figure S1. Functional connectivity depends on smoothing kernel.** sgACC functional connectivity depends on the whole-grey-matter smoothing kernel. Bilateral sgACC FC was extracted with 10-mm seed and target radii, grey-matter constraints and GSR, using unsmoothed mean seed time series and median target FC. The 80 participant–target combinations contributing to the primary stimulated-target analyses provide 320 values across four kernels (0, 3, 6 and 12 mm FWHM). Sample sizes were 8, 14, 15, 13, 15 and 15 for 5 cm, F3, F3 anterior, F3 lateral, F3 medial and F3 posterior, respectively. The main analysis uses 3-mm whole-GM FWHM.

#### Generalized additive models

Because HBC is known to show nonlinear behavior [12], we additionally fit generalized additive models (GAMs) with the mgcv package. For the stimulated-target dataset, the GAM was specified as follows:

$$\text{model\_gam}_{\text{HBC\sim ExFC}} = \Delta HBC \sim$$

### parametric terms

Session + *TargetCortex* +

### smooth terms

s(FC, k = 50) + s(E, k = 20) + s(Pain, k = 6) + *SideEffects* + s(PCD, k = 50) +

### interaction terms

s(E, by = *TargetCortex*) + s(FC, by = *TargetCortex*, k = 80) + ti(E, FC) + ti(E, Pain, k = 6) + ti(E, FC, by = *TargetCortex*) + ti(PCD, E) + s(PCD, by = *TargetCortex*, k = 30) +

### random terms

s(SubID, by = Session, bs = "re") + s(SubID, by = E, bs = "fs")

Parametric terms (Session, TargetCortex and SideEffects) were included, while smooth terms for FC, E, Pain and PCD and their interactions allowed nonlinear relationships. Participant-level variation was represented by subject-by-session random-effect terms (s(SubID, by = Session, bs = "re")) and an E-field-dependent subject random-effect term (s(SubID, by = E, bs = "fs")).

In the GAM formulation, s() denotes univariate smooth functions of predictors (thin plate regression splines by default), and ti() denotes tensor-product interaction smooths that model nonlinear interactions in an ANOVA-like decomposition (main-effect smooths plus interaction smooths). The argument k specifies the basis dimension, i.e., the size of the basis used to represent a smooth (which sets an upper bound on its effective degrees of freedom). For tensor-product interactions, k specifies the dimensions of the marginal bases. The actual smoothness is determined by penalization during fitting, so k acts as a flexible complexity cap rather than directly fixing the wiggliness. Basis dimensions were retained from the original model specification, while smoothing parameters were estimated by fast restricted maximum likelihood using mgcv::bam (v1.9.1; method = "fREML", discrete = TRUE).

#### GAM reference comparison

**Table S1.** Fit of E-only and full generalized additive models.

| Dataset | Model | Rows | AIC | ΔAIC vs E-only | Adjusted $R^2$ |
|---|---|---|---|---|---|
| Stimulated target | E-field only | 3 502 | 12812.56 | 0.00 | 0.1939 |
| Stimulated target | E-field + FC | 3 502 | 12313.35 | -499.21 | 0.3163 |
| Six locations | E-field only | 21 012 | 77316.85 | 0.00 | 0.1559 |
| Six locations | E-field + FC | 21 012 | 77166.31 | -150.54 | 0.1668 |

**Notes.** All models use GSR, 10-mm seed/target radii and 3-mm whole-GM FWHM in 15 participants. Full models use bilateral sgACC, mean seed time series and median target FC. The E-only GAM removes s(FC), target-specific s(FC), ti(E,FC) and target-specific ti(E,FC). Remaining terms and basis dimensions are unchanged. Fits use fREML with discrete estimation. AIC derives from log likelihood and effective parameter count. ΔAIC is relative to E-only within each dataset. Six-location scores describe the expanded dataset.

**Models with and without covariates**

Using the same 3,502 observations and original observation selection, we refitted the stimulated-target models after omitting pain, side effects, PCD and all associated interactions. Session, target, participant random effects, core dose/FC terms and GAM basis dimensions were retained. LMM comparisons used ML, with REML estimates for $R^2$. In the LMMs, E-field terms and FC terms were each tested jointly with their respective interactions. GAM comparisons used AIC and adjusted $R^2$ (Table S2).

In the LMM without these covariate terms, adding E-field and its target interactions improved fit relative to Session, target and participant random effects alone ($\chi^2(6) = 122.65$, $p = 4.51 \times 10^{-24}$).

**Table S2. Sensitivity to covariate adjustment.**

**A. E-field and FC contributions**

| Comparison | With covariates | Without covariates |
|---|---|---|
| E-field vs. MT, LMM AIC decrease | 45.61 | 43.14 |
| Adding FC, LMM AIC decrease | 54.42 | 64.26 |
| Adding FC, LMM $\chi^2$ (df) | 78.419 (12) | 88.261 (12) |
| Adding FC, LMM p | $8.257 \times 10^{-12}$ | $1.071 \times 10^{-13}$ |
| Adding FC, GAM AIC decrease | 499.21 | 594.93 |

**B. LMM fit indices**

| Model | With covariates | | | Without covariates | | |
|---|---|---|---|---|---|---|
| | AIC | $R^2$ (cond.) | $R^2$ (marg.) | AIC | $R^2$ (cond.) | $R^2$ (marg.) |
| $MT$ | 13047.76 | 0.168 | 0.061 | 13156.66 | 0.129 | 0.025 |
| $MT \times FC$ | 13005.06 | 0.188 | 0.077 | 13110.45 | 0.153 | 0.044 |
| $E$ | 13002.15 | 0.206 | 0.085 | 13113.52 | 0.163 | 0.048 |
| $E \times FC$ | **12947.73** | **0.236** | **0.104** | **13049.26** | **0.202** | **0.073** |

**C. GAM fit indices**

| Model | With covariates | | Without covariates | |
|---|---|---|---|---|
| | AIC | Adjusted $R^2$ | AIC | Adjusted $R^2$ |
| $E$ | 12812.56 | 0.194 | 13005.18 | 0.140 |
| $E \times FC$ | **12313.35** | **0.316** | **12410.25** | **0.288** |

**Notes.** Covariates denote pain, side effects and PCD and their associated interactions. In panel A, AIC decrease is AIC before minus AIC after using E-field or adding FC. The FC comparisons include all FC terms and interactions. LMM AIC and likelihood-ratio tests use ML, and $R^2$ uses REML. GAMs use fREML. P values are nominal. AIC differences use unrounded values. All models use bilateral sgACC, mean seed time series, median target FC, 10-mm seed/target radii, 3-mm FWHM and GSR. Bold marks the best fit indices within each covariate set.

#### Control-seed analyses

We refitted the original full GAMs and LMMs after replacing sgACC–target FC with FC between the same target locations and each control seed. Each model used the same recorded observations, covariates, 10-mm seed and target radii, 3-mm whole-GM FWHM and GSR. Seed time series were averaged across voxels, and target FC was the median of non-zero projected Fisher-z values. AIC differences were calculated relative to bilateral sgACC within each dataset (Table S3), with single-right sgACC comparisons in Table S4. The stimulated-target models are primary, and six-location fits are descriptive extensions.

The bilateral sgACC seed combined spheres centred at MNI (−6, 16, −10) and (6, 16, −10) as a union in native space before time-series averaging. The single-right reference used MNI (6, 16, −10) (Fox et al. [39]). Control centres followed Van Dijk et al. [51] for PCC (0, −53, 26), Vincent et al. [52] (Table 2) for left/right posterior IPL (−47, −71, 29 and 50, −64, 27), and Philippi et al. [53] (Table 2) for left/right visual cortex (−19, −98, 3 and 19, −98, 3). All seeds were grey-matter constrained. The same seed-mean, Fisher transformation, surface projection and target-median extraction order was applied to all seeds.

**Table S3.** Control-seed model comparisons with bilateral sgACC and GSR.

| Dataset | Seed | GAM AIC | GAM ΔAIC | GAM adjusted $R^2$ | LMM AIC | LMM ΔAIC | LMM $R^2m$ / $R^2c$ |
|---|---|---|---|---|---|---|---|
| Target | Bilateral sgACC | 12313.35 | 0.00 | 0.3163 | 12947.73 | 0.00 | 0.1044 / 0.2362 |
| Target | Left IPL | 12434.53 | 121.17 | 0.2923 | 12989.83 | 42.10 | 0.0943 / 0.2146 |
| Target | Right IPL | 12358.06 | 44.71 | 0.3058 | 12976.17 | 28.44 | 0.0962 / 0.2083 |
| Target | Left visual cortex | 12392.60 | 79.25 | 0.3004 | 12964.44 | 16.71 | 0.1010 / 0.2209 |
| Target | Right visual cortex | 12327.26 | 13.91 | 0.3150 | 12958.42 | 10.69 | 0.1079 / 0.2400 |
| Target | PCC | 12364.33 | 50.98 | 0.3052 | 12982.48 | 34.75 | 0.1038 / 0.2358 |
| Six locations | Bilateral sgACC | 77166.31 | 0.00 | 0.1668 | 78275.42 | 0.00 | 0.0254 / 0.1373 |
| Six locations | Left IPL | 77278.27 | 111.96 | 0.1603 | 78258.76 | -16.65 | 0.0261 / 0.1370 |
| Six locations | Right IPL | 77253.89 | 87.58 | 0.1615 | 78258.18 | -17.24 | 0.0267 / 0.1385 |
| Six locations | Left visual cortex | 77249.02 | 82.71 | 0.1628 | 78276.05 | 0.63 | 0.0256 / 0.1368 |
| Six locations | Right visual cortex | 77196.43 | 30.12 | 0.1646 | 78277.47 | 2.05 | 0.0257 / 0.1369 |
| Six locations | PCC | 77273.23 | 106.92 | 0.1607 | 78278.04 | 2.62 | 0.0254 / 0.1357 |

***Notes.*** ΔAIC = control AIC − bilateral sgACC AIC within each dataset. Positive values favour sgACC. GAMs use fREML. LMM AIC uses ML, stimulated-target $R^2$ uses REML, and six-location $R^2$ uses ML. $R^2m$/$R^2c$ denote marginal/conditional $R^2$. Six-location fits are descriptive.

**Table S4.** Control-seed model comparisons with single-right sgACC and GSR.

| Dataset | Seed | GAM AIC | GAM ΔAIC | GAM adjusted $R^2$ | LMM AIC | LMM ΔAIC | LMM $R^2m$ / $R^2c$ |
|---|---|---|---|---|---|---|---|
| Target | Single-right sgACC | 12306.97 | 0.00 | 0.3177 | 12951.70 | 0.00 | 0.1037 / 0.2334 |

| Dataset | Seed | GAM AIC | GAM ΔAIC | GAM adjusted $R^2$ | LMM AIC | LMM ΔAIC | LMM $R^2m$ / $R^2c$ |
|---|---|---|---|---|---|---|---|
| Target | Left IPL | 12434.53 | 127.55 | 0.2923 | 12989.83 | 38.13 | 0.0943 / 0.2146 |
| Target | Right IPL | 12358.06 | 51.09 | 0.3058 | 12976.17 | 24.47 | 0.0962 / 0.2083 |
| Target | Left visual cortex | 12392.60 | 85.62 | 0.3004 | 12964.44 | 12.74 | 0.1010 / 0.2209 |
| Target | Right visual cortex | 12327.26 | 20.29 | 0.3150 | 12958.42 | 6.72 | 0.1079 / 0.2400 |
| Target | PCC | 12364.33 | 57.36 | 0.3052 | 12982.48 | 30.78 | 0.1038 / 0.2358 |
| Six locations | Single-right sgACC | 77236.49 | 0.00 | 0.1625 | 78269.13 | 0.00 | 0.0258 / 0.1378 |
| Six locations | Left IPL | 77278.27 | 41.77 | 0.1603 | 78258.76 | -10.37 | 0.0261 / 0.1370 |
| Six locations | Right IPL | 77253.89 | 17.40 | 0.1615 | 78258.18 | -10.95 | 0.0267 / 0.1385 |
| Six locations | Left visual cortex | 77249.02 | 12.52 | 0.1628 | 78276.05 | 6.92 | 0.0256 / 0.1368 |
| Six locations | Right visual cortex | 77196.43 | -40.07 | 0.1646 | 78277.47 | 8.33 | 0.0257 / 0.1369 |
| Six locations | PCC | 77273.23 | 36.74 | 0.1607 | 78278.04 | 8.90 | 0.0254 / 0.1357 |

***Notes.*** ΔAIC = control AIC − single-right sgACC AIC within each dataset. Positive values favour sgACC. GAMs use fREML. LMM AIC uses ML, stimulated-target $R^2$ uses REML, and six-location $R^2$ uses ML. $R^2m$/$R^2c$ denote marginal/conditional $R^2$. Six-location fits are descriptive.

#### Linear mixed effects models

##### Model comparison: MSO versus cortical E-field.

Linear mixed-effects models were specified as described in the main Methods ($model_{HBC\sim MT}$, $model_{HBC\sim E}$). Both models included random intercepts and session slopes per subject and the same covariates (*Session, pain, side effects, pulsewise coil displacement,* and their interactions with dose and target), differing only in the primary dosing term (%MT vs. cortical E-field). The E-field model had lower ML-based information criteria (AIC, AICc and BIC) and higher marginal and conditional $R^2$ than the %MT model (Table 1). Coefficient estimates and fit summaries are reported in Table S5. Adjusted ICC (0.132 versus 0.113) and conditional training RMSE (1.503 versus 1.514) were similar. Residuals and random effects were inspected visually.

**Table S5.** Relative motor-threshold dose (%MT) versus cortical E-field strength.

| | **ΔHBC ~ MT** | | **ΔHBC ~ E-field** | |
|---|---|---|---|---|
| ***Predictors*** | ***Estimates*** | ***p*** | ***Estimates*** | ***p*** |
| Intercept | 0.30 | 0.050 | 0.28 | 0.082 |
| Session [2] | -0.04 | 0.833 | -0.05 | 0.784 |
| Session [3] | -0.07 | 0.694 | -0.10 | 0.588 |
| TMS condition \| target cortex | | | | |
| F3 | -0.09 | 0.422 | -0.06 | 0.577 |
| F3 ant | 0.11 | 0.344 | 0.22 | 0.058 |
| F3 lat | 0.26 | **0.026** | 0.33 | **0.005** |
| F3 med | -0.06 | 0.595 | -0.02 | 0.841 |
| F3 pos | 0.05 | 0.663 | -0.02 | 0.879 |
| %MT \| E | 0.01 | **<0.001** | 0.01 | **<0.001** |
| Pain | 0.03 | 0.491 | -0.01 | 0.854 |
| Side effects | -0.16 | **<0.001** | -0.22 | **<0.001** |
| PCD | -0.01 | 0.858 | -0.00 | 0.947 |
| TMS condition × %MT \| target cortex × E | | | | |
| %MT \| E × F3 | 0.00 | 0.931 | 0.01 | 0.060 |
| %MT \| E × F3 ant | -0.00 | 0.339 | -0.00 | 0.511 |
| %MT \| E × F3 lat | 0.00 | 0.531 | 0.01 | 0.058 |
| %MT \| E × F3 med | -0.00 | 0.682 | -0.01 | 0.064 |
| %MT \| E × F3 pos | 0.00 | 0.896 | 0.01 | 0.239 |
| PCD × %MT \| E | 0.00 | 0.450 | 0.00 | 0.233 |
| PCD × TMS condition \| target cortex | | | | |
| PCD × F3 | 0.56 | **0.004** | 0.48 | **0.013** |
| PCD × F3 ant | 0.46 | **0.014** | 0.47 | **0.012** |
| PCD × F3 lat | 0.47 | **0.014** | 0.39 | **0.046** |
| PCD × F3 med | 0.08 | 0.638 | 0.12 | 0.506 |
| PCD × F3 pos | 1.15 | **<0.001** | 1.10 | **<0.001** |
| Pain × %MT \| E | 0.00 | **<0.001** | 0.00 | **0.005** |
| ICC | 0.11 | | 0.13 | |
| N | 15 SubID | | 15 SubID | |
| Observations | 3502 | | 3502 | |
| Marg. $R^2$ / Cond. $R^2$ | 0.061 / 0.168 | | 0.085 / 0.206 | |
| AIC | 13047.757 | | 13002.148 | |

***Notes.*** Coefficients, ICC and $R^2$ in Table S5 are from REML fits. AIC is from matched ML fits, as in Table 1. The ΔHBC ~ MT model uses stimulator output intensity (maximum stimulator output, MSO) expressed as percentage of the individual motor threshold (*%MT*) for each coil placement (*coil condition*) to predict change in heart-brain-coupling (*HBC*). In contrast, the ΔHBC ~ E-field model uses cortical stimulation strength |E| in V/m (*E*), extracted in grey matter under each coil placement (*target cortex*).

#### FC-extended E-field models.

Adding FC and its interactions to the E-field LMM ($model_{HBC \sim E \times FC}$) improved its ML fit (likelihood-ratio $\chi^2(12) = 78.42$, P = $8.26 \times 10^{-12}$), with AIC decreasing from 13002.15 to 12947.73. REML marginal $R^2$ increased from 0.085 to 0.104 and conditional $R^2$ from 0.206 to 0.236. The parallel %MT comparison yielded $\chi^2(12) = 66.70$, P = $1.32 \times 10^{-9}$. These P values are nominal and unadjusted. Tables S6 and S7 report the %MT and E-field comparisons, respectively; Table S8 compares the stimulated-target and six-extraction-site analyses.

**Table S6.** Relative motor-threshold dose (%MT): effects of functional connectivity.

| | ΔHBC ~ MT | | | ΔHBC ~ MT + FC | | |
|---|---|---|---|---|---|---|
| ***Predictors*** | ***Estimates*** | ***CI*** | ***p*** | ***Estimates*** | ***CI*** | ***p*** |
| Intercept | 0.30 | -0.00 – 0.61 | 0.050 | 0.30 | -0.03 – 0.63 | 0.074 |
| Session [2] | -0.04 | -0.46 – 0.37 | 0.833 | -0.04 | -0.45 – 0.38 | 0.846 |
| Session [3] | -0.07 | -0.46 – 0.31 | 0.694 | -0.07 | -0.45 – 0.31 | 0.703 |
| TMS condition | | | | | | |
| F3 | -0.09 | -0.30 – 0.13 | 0.422 | -0.14 | -0.38 – 0.09 | 0.236 |
| F3 ant | 0.11 | -0.11 – 0.33 | 0.344 | 0.08 | -0.16 – 0.31 | 0.522 |
| F3 lat | 0.26 | 0.03 – 0.49 | **0.026** | 0.14 | -0.10 – 0.38 | 0.245 |
| F3 med | -0.06 | -0.26 – 0.15 | 0.595 | -0.01 | -0.23 – 0.22 | 0.946 |
| F3 pos | 0.05 | -0.16 – 0.25 | 0.663 | 0.01 | -0.21 – 0.23 | 0.928 |
| %MT | 0.01 | 0.01 – 0.01 | **<0.001** | 0.01 | 0.01 – 0.01 | **<0.001** |
| Pain | 0.03 | -0.06 – 0.12 | 0.491 | 0.06 | -0.04 – 0.15 | 0.218 |
| Side effects | -0.16 | -0.24 – -0.07 | **<0.001** | -0.16 | -0.24 – -0.07 | **<0.001** |
| PCD | -0.01 | -0.10 – 0.08 | 0.858 | -0.01 | -0.10 – 0.08 | 0.875 |
| %MT × TMS condition | | | | | | |
| F3 × %MT | 0.00 | -0.01 – 0.01 | 0.931 | -0.00 | -0.01 – 0.01 | 0.525 |
| F3 ant × %MT | -0.00 | -0.01 – 0.01 | 0.339 | -0.01 | -0.02 – 0.00 | 0.234 |
| F3 lat × %MT | 0.00 | -0.01 – 0.01 | 0.531 | 0.00 | -0.01 – 0.01 | 0.830 |
| F3 med × %MT | -0.00 | -0.01 – 0.01 | 0.682 | -0.01 | -0.02 – 0.00 | 0.296 |
| F3 pos × %MT | 0.00 | -0.01 – 0.01 | 0.896 | 0.00 | -0.01 – 0.01 | 0.873 |
| %MT × PCD | 0.00 | -0.00 – 0.01 | 0.450 | 0.00 | -0.00 – 0.00 | 0.532 |
| PCD × TMS condition | | | | | | |
| PCD × F3 | 0.56 | 0.18 – 0.93 | **0.004** | 0.54 | 0.17 – 0.92 | **0.005** |
| PCD × F3 ant | 0.46 | 0.10 – 0.83 | **0.014** | 0.48 | 0.11 – 0.85 | **0.011** |
| PCD × F3 lat | 0.47 | 0.10 – 0.85 | **0.014** | 0.52 | 0.15 – 0.90 | **0.007** |
| PCD × F3 med | 0.08 | -0.27 – 0.43 | 0.638 | 0.09 | -0.26 – 0.44 | 0.603 |
| PCD × F3 pos | 1.15 | 0.79 – 1.51 | **<0.001** | 1.12 | 0.76 – 1.48 | **<0.001** |
| %MT × Pain | 0.00 | 0.00 – 0.01 | **<0.001** | 0.00 | 0.00 – 0.01 | **<0.001** |
| FC | | | | -0.90 | -1.74 – -0.05 | **0.037** |
| FC × TMS condition | | | | | | |
| FC × F3 | | | | -0.18 | -3.09 – 2.73 | 0.903 |
| FC × F3 ant | | | | 0.89 | -1.99 – 3.77 | 0.546 |
| FC × F3 lat | | | | -5.09 | -8.00 – -2.19 | **<0.001** |
| FC × F3 med | | | | -0.59 | -3.19 – 2.01 | 0.657 |
| FC × F3 pos | | | | 2.95 | 0.11 – 5.79 | **0.042** |
| FC × %MT | | | | -0.01 | -0.04 – 0.02 | 0.489 |
| FC × %MT × TMS condition | | | | | | |
| FC × F3 × %MT | | | | -0.08 | -0.21 – 0.05 | 0.204 |
| FC × F3 ant × %MT | | | | -0.03 | -0.16 – 0.10 | 0.614 |
| FC × F3 lat × %MT | | | | -0.05 | -0.18 – 0.08 | 0.477 |
| FC × F3 med × %MT | | | | 0.06 | -0.07 – 0.18 | 0.371 |
| FC × F3 pos × %MT | | | | 0.03 | -0.11 – 0.17 | 0.655 |
| ICC | | 0.11 | | | 0.12 | |
| N | | 15 SubID | | | 15 SubID | |
| Observations | | 3502 | | | 3502 | |
| Marg. $R^2$ / Cond. $R^2$ | | 0.061 / 0.168 | | | 0.077 / 0.188 | |
| AIC | | 13047.757 | | | 13005.061 | |
| AICc | | 13048.329 | | | 13006.155 | |

***Notes.*** Coefficients, ICC and $R^2$ in this table are from REML fits. Information criteria are from matched ML fits, as in Table 1. The bilateral sgACC seed was the native-space union of grey-matter-constrained, 10-mm-radius spheres centred at MNI (−6, 16, −10) and (6, 16, −10). Its unsmoothed mean time series was correlated with whole-grey-matter time series smoothed at 3-mm FWHM, with GSR. Correlations were Fisher transformed and projected to native cortical elements with the corrected three-axis sampler, then summarized by their median within 10-mm grey-matter target spheres.

**Table S7.** Cortical stimulation strength (E): Effects of functional connectivity.

| | ΔHBC ~ E-field | | | ΔHBC ~ E-field + FC | | |
|---|---|---|---|---|---|---|
| ***Predictors*** | ***Estimates*** | ***CI*** | ***p*** | ***Estimates*** | ***CI*** | ***p*** |
| Intercept | 0.28 | -0.04 – 0.61 | 0.082 | 0.28 | -0.08 – 0.64 | 0.114 |
| Session [2] | -0.05 | -0.47 – 0.36 | 0.784 | -0.05 | -0.46 – 0.37 | 0.810 |
| Session [3] | -0.10 | -0.47 – 0.28 | 0.588 | -0.09 | -0.47 – 0.28 | 0.606 |
| Target cortex | | | | | | |
| F3 | -0.06 | -0.28 – 0.15 | 0.577 | -0.16 | -0.41 – 0.09 | 0.201 |
| F3 ant | 0.22 | -0.01 – 0.44 | 0.058 | 0.16 | -0.09 – 0.41 | 0.217 |
| F3 lat | 0.33 | 0.10 – 0.56 | **0.005** | 0.16 | -0.10 – 0.41 | 0.224 |
| F3 med | -0.02 | -0.23 – 0.18 | 0.841 | -0.03 | -0.27 – 0.21 | 0.787 |
| F3 pos | -0.02 | -0.22 – 0.19 | 0.879 | -0.10 | -0.33 – 0.14 | 0.417 |
| E | 0.01 | 0.01 – 0.02 | **<0.001** | 0.01 | 0.01 – 0.01 | **<0.001** |
| Pain | -0.01 | -0.10 – 0.08 | 0.854 | -0.01 | -0.11 – 0.09 | 0.846 |
| Side effects | -0.22 | -0.31 – -0.13 | **<0.001** | -0.21 | -0.30 – -0.12 | **<0.001** |
| PCD | -0.00 | -0.09 – 0.09 | 0.947 | 0.01 | -0.08 – 0.09 | 0.909 |
| E × Target cortex | | | | | | |
| E × F3 | 0.01 | -0.00 – 0.02 | 0.060 | 0.00 | -0.01 – 0.01 | 0.917 |
| E × F3 ant | -0.00 | -0.01 – 0.01 | 0.511 | -0.01 | -0.02 – 0.01 | 0.324 |
| E × F3 lat | 0.01 | -0.00 – 0.02 | 0.058 | 0.01 | -0.00 – 0.02 | 0.229 |
| E × F3 med | -0.01 | -0.02 – 0.00 | 0.064 | -0.01 | -0.02 – -0.00 | **0.023** |
| E × F3 pos | 0.01 | -0.00 – 0.02 | 0.239 | 0.01 | -0.00 – 0.02 | 0.244 |
| E × PCD | 0.00 | -0.00 – 0.01 | 0.233 | 0.00 | -0.00 – 0.00 | 0.324 |
| PCD × Target cortex | | | | | | |
| PCD × F3 | 0.48 | 0.10 – 0.86 | **0.013** | 0.41 | 0.02 – 0.79 | **0.038** |
| PCD × F3 ant | 0.47 | 0.10 – 0.84 | **0.012** | 0.47 | 0.09 – 0.84 | **0.014** |
| PCD × F3 lat | 0.39 | 0.01 – 0.77 | **0.046** | 0.42 | 0.03 – 0.80 | **0.034** |
| PCD × F3 med | 0.12 | -0.23 – 0.47 | 0.506 | 0.09 | -0.27 – 0.44 | 0.636 |
| PCD × F3 pos | 1.10 | 0.74 – 1.46 | **<0.001** | 1.04 | 0.68 – 1.41 | **<0.001** |
| E × Pain | 0.00 | 0.00 – 0.01 | **0.005** | 0.00 | 0.00 – 0.01 | **0.006** |
| FC | | | | -1.14 | -2.01 – -0.27 | **0.010** |
| FC × Target cortex | | | | | | |
| FC × F3 | | | | 0.01 | -3.04 – 3.07 | 0.993 |
| FC × F3 ant | | | | 1.70 | -1.33 – 4.73 | 0.272 |
| FC × F3 lat | | | | -4.87 | -7.94 – -1.80 | **0.002** |
| FC × F3 med | | | | 0.16 | -2.60 – 2.91 | 0.910 |
| FC × F3 pos | | | | 2.81 | -0.22 – 5.83 | 0.069 |
| FC × E | | | | -0.02 | -0.06 – 0.01 | 0.181 |
| FC × E × Target cortex | | | | | | |
| FC × E × F3 | | | | -0.23 | -0.37 – -0.08 | **0.002** |
| FC × E × F3 ant | | | | -0.11 | -0.26 – 0.03 | 0.128 |
| FC × E × F3 lat | | | | -0.15 | -0.29 – -0.00 | **0.043** |
| FC × E × F3 med | | | | 0.01 | -0.13 – 0.15 | 0.934 |
| FC × E × F3 pos | | | | -0.04 | -0.18 – 0.10 | 0.564 |
| ICC | | 0.13 | | | 0.15 | |
| N | | 15 SubID | | | 15 SubID | |
| Observations | | 3502 | | | 3502 | |
| Marg. $R^2$ / Cond. $R^2$ | | 0.085 / 0.206 | | | 0.104 / 0.236 | |
| AIC | | 13002.148 | | | 12947.729 | |

***Notes.*** Coefficients, ICC and $R^2$ are from REML fits. Information criteria are from matched ML fits, as in Table 1. FC processing follows Table S6. Cortical dose was the 99th percentile of |E| in the grey-matter target sphere. Tables S9–S12 vary the sgACC seed radius (labelled FC, 5 or 10 mm), the shared E-field/target-FC extraction radius (labelled E, 5 or 10 mm), and whole-grey-matter FWHM (0, 3, 6 or 12 mm). Grid coefficients, ICC and $R^2$ use REML, and information criteria use matched ML fits. Tables S3 and S4 compare controls with bilateral and single-right sgACC, respectively, using GSR. Table S1 compares the E-only and full bilateral-sgACC GAMs within each dataset.

**Table S8.** Stimulated-target versus six-extraction-site LMMs.

| | ΔHBC ~ E + FC<br>TMS condition == target cortex | | | ΔHBC ~ E + FC<br>all data | | |
|---|---|---|---|---|---|---|
| ***Predictors*** | ***Estimates*** | ***CI*** | ***p*** | ***Estimates*** | ***CI*** | ***p*** |
| Intercept | 0.28 | -0.08 – 0.64 | 0.114 | 0.27 | — | — |
| Session [2] | -0.05 | -0.46 – 0.37 | 0.810 | -0.02 | — | — |
| Session [3] | -0.09 | -0.47 – 0.28 | 0.606 | -0.10 | — | — |
| Target cortex | | | | | | |
| F3 | -0.16 | -0.41 – 0.09 | 0.201 | -0.08 | — | — |
| F3 ant | 0.16 | -0.09 – 0.41 | 0.217 | 0.03 | — | — |
| F3 lat | 0.16 | -0.10 – 0.41 | 0.224 | -0.02 | — | — |
| F3 med | -0.03 | -0.27 – 0.21 | 0.787 | 0.00 | — | — |
| F3 pos | -0.10 | -0.33 – 0.14 | 0.417 | -0.02 | — | — |
| FC | -1.14 | -2.01 – -0.27 | **0.010** | -0.10 | — | — |
| E | 0.01 | 0.01 – 0.01 | **<0.001** | 0.01 | — | — |
| Pain | -0.01 | -0.11 – 0.09 | 0.846 | 0.10 | — | — |
| Side effects | -0.21 | -0.30 – -0.12 | **<0.001** | -0.16 | — | — |
| PCD | 0.01 | -0.08 – 0.09 | 0.909 | 0.08 | — | — |
| E × Target cortex | | | | | | |
| E × F3 | 0.00 | -0.01 – 0.01 | 0.917 | 0.00 | — | — |
| E × F3 ant | -0.01 | -0.02 – 0.01 | 0.324 | 0.00 | — | — |
| E × F3 lat | 0.01 | -0.00 – 0.02 | 0.229 | 0.00 | — | — |
| E × F3 med | -0.01 | -0.02 – -0.00 | **0.023** | 0.00 | — | — |
| E × F3 pos | 0.01 | -0.00 – 0.02 | 0.244 | 0.00 | — | — |
| FC × Target cortex | | | | | | |
| FC × F3 | 0.01 | -3.04 – 3.07 | 0.993 | 0.10 | — | — |
| FC × F3 ant | 1.70 | -1.33 – 4.73 | 0.272 | -0.10 | — | — |
| FC × F3 lat | -4.87 | -7.94 – -1.80 | **0.002** | 0.19 | — | — |
| FC × F3 med | 0.16 | -2.60 – 2.91 | 0.910 | 0.04 | — | — |
| FC × F3 pos | 2.81 | -0.22 – 5.83 | 0.069 | 0.22 | — | — |
| E × PCD | 0.00 | -0.00 – 0.00 | 0.324 | 0.00 | — | — |
| PCD × Target cortex | | | | | | |
| PCD × F3 | 0.41 | 0.02 – 0.79 | **0.038** | -0.06 | — | — |
| PCD × F3 ant | 0.47 | 0.09 – 0.84 | **0.014** | -0.02 | — | — |
| PCD × F3 lat | 0.42 | 0.03 – 0.80 | **0.034** | -0.06 | — | — |
| PCD × F3 med | 0.09 | -0.27 – 0.44 | 0.636 | -0.00 | — | — |
| PCD × F3 pos | 1.04 | 0.68 – 1.41 | **<0.001** | -0.02 | — | — |
| FC × E | -0.02 | -0.06 – 0.01 | 0.181 | -0.01 | — | — |
| E × Pain | 0.00 | 0.00 – 0.01 | **0.006** | 0.00 | — | — |
| FC × E × Target cortex | | | | | | |
| FC × E × F3 | -0.23 | -0.37 – -0.08 | **0.002** | -0.02 | — | — |
| FC × E × F3 ant | -0.11 | -0.26 – 0.03 | 0.128 | -0.06 | — | — |
| FC × E × F3 lat | -0.15 | -0.29 – -0.00 | **0.043** | -0.06 | — | — |
| FC × E × F3 med | 0.01 | -0.13 – 0.15 | 0.934 | -0.01 | — | — |
| FC × E × F3 pos | -0.04 | -0.18 – 0.10 | 0.564 | -0.02 | — | — |
| ICC | 0.15 | | | 0.11 | | |
| N | 15 SubID | | | 15 SubID | | |
| Observations | 3502 | | | 21012 | | |

***Notes.*** The stimulated-*target* analysis pairs each response with FC and E-field at the cortical location corresponding to its coil condition (3,502 observations). The descriptive six-extraction-site extension pairs each response with FC and E-field at all six extraction locations (21,012 rows). Stimulated-target coefficients use REML and six-location coefficients use ML. Both include 15 participants and exclude 315 copied 5-cm stimulation observations; 357 recorded 5-cm observations from eight participants remain. The six-site coefficients are descriptive; their CI and P entries are not reported (—).

**Table S9.** Effects of extraction sphere size and smoothing (E = 5 mm, FC = 5 mm).

| | ΔHBC ~ E + FC | | ΔHBC ~ E + FC | | ΔHBC ~ E + FC | | ΔHBC ~ E + FC | |
|---|---|---|---|---|---|---|---|---|
| **ROI E** | **5 mm** | | **5 mm** | | **5 mm** | | **5 mm** | |
| **ROI FC** | **5 mm** | | **5 mm** | | **5 mm** | | **5 mm** | |
| **FC smoothing** | **0 mm** | | **3 mm** | | **6 mm** | | **12 mm** | |
| ***Predictors*** | ***Estimates*** | ***p*** | ***Estimates*** | ***p*** | ***Estimates*** | ***p*** | ***Estimates*** | ***p*** |
| Intercept | 0.28 | 0.082 | 0.27 | 0.095 | 0.25 | 0.110 | 0.26 | 0.098 |
| Session [2] | -0.06 | 0.766 | -0.05 | 0.795 | -0.05 | 0.819 | -0.05 | 0.811 |
| Session [3] | -0.11 | 0.546 | -0.10 | 0.582 | -0.09 | 0.614 | -0.09 | 0.611 |
| Target cortex | | | | | | | | |
| F3 | 0.11 | 0.348 | 0.05 | 0.668 | 0.07 | 0.589 | 0.14 | 0.270 |
| F3 ant | 0.47 | **<0.001** | 0.32 | **0.012** | 0.31 | **0.015** | 0.38 | **0.003** |
| F3 lat | 0.52 | **<0.001** | 0.38 | **0.002** | 0.39 | **0.002** | 0.47 | **<0.001** |
| F3 med | 0.06 | 0.608 | 0.01 | 0.910 | 0.03 | 0.820 | 0.07 | 0.540 |
| F3 pos | -0.01 | 0.933 | -0.04 | 0.732 | 0.00 | 0.980 | 0.09 | 0.446 |
| FC | 1.23 | 0.154 | -0.02 | 0.957 | 0.24 | 0.446 | 0.39 | 0.150 |
| E | 0.01 | **<0.001** | 0.01 | **<0.001** | 0.01 | **<0.001** | 0.01 | **<0.001** |
| Pain | -0.01 | 0.812 | -0.00 | 0.936 | -0.01 | 0.791 | -0.04 | 0.452 |
| Side effects | -0.25 | **<0.001** | -0.22 | **<0.001** | -0.19 | **<0.001** | -0.20 | **<0.001** |
| PCD | 0.01 | 0.767 | 0.01 | 0.828 | 0.01 | 0.811 | 0.01 | 0.832 |
| E × Target cortex | | | | | | | | |
| E × F3 | 0.01 | 0.061 | 0.01 | 0.303 | 0.00 | 0.909 | -0.00 | 0.662 |
| F3 ant ×E | 0.00 | 0.859 | -0.00 | 0.485 | -0.01 | 0.247 | -0.01 | 0.250 |
| F3 lat ×E | 0.01 | 0.207 | 0.00 | 0.455 | 0.00 | 0.501 | 0.01 | 0.294 |
| F3 med ×E | -0.01 | **0.029** | -0.01 | **0.013** | -0.01 | **0.007** | -0.01 | **0.014** |
| E × F3 pos | 0.01 | 0.293 | 0.00 | 0.375 | 0.01 | 0.354 | 0.00 | 0.471 |
| FC × Target cortex | | | | | | | | |
| FC × F3 | -10.30 | **0.002** | -4.13 | **0.005** | -3.37 | **<0.001** | -2.52 | **0.004** |
| FC × F3 ant | -2.65 | 0.446 | -2.40 | 0.114 | -2.97 | **0.003** | -2.52 | **0.004** |
| FC × F3 lat | -17.69 | **<0.001** | -7.12 | **<0.001** | -5.29 | **<0.001** | -4.47 | **<0.001** |
| FC × F3 med | -6.82 | **0.032** | -2.47 | 0.078 | -1.92 | **0.049** | -1.66 | **0.044** |
| FC × F3 pos | -7.83 | **0.013** | -3.16 | **0.030** | -1.93 | 0.052 | -1.62 | 0.061 |
| E × PCD | 0.00 | 0.307 | 0.00 | 0.297 | 0.00 | 0.275 | 0.00 | 0.352 |
| PCD × Target cortex | | | | | | | | |
| PCD × F3 | 0.42 | **0.033** | 0.41 | **0.042** | 0.35 | 0.080 | 0.37 | 0.063 |
| PCD × F3 ant | 0.44 | **0.024** | 0.44 | **0.024** | 0.39 | **0.045** | 0.41 | **0.033** |
| PCD × F3 lat | 0.39 | **0.050** | 0.39 | 0.050 | 0.34 | 0.083 | 0.37 | 0.064 |

| | ΔHBC ~ E + FC | | ΔHBC ~ E + FC | | ΔHBC ~ E + FC | | ΔHBC ~ E + FC | |
|---|---|---|---|---|---|---|---|---|
| **ROI E** | **5 mm** | | **5 mm** | | **5 mm** | | **5 mm** | |
| **ROI FC** | **5 mm** | | **5 mm** | | **5 mm** | | **5 mm** | |
| **FC smoothing** | **0 mm** | | **3 mm** | | **6 mm** | | **12 mm** | |
| ***Predictors*** | ***Estimates*** | ***p*** | ***Estimates*** | ***p*** | ***Estimates*** | ***p*** | ***Estimates*** | ***p*** |
| PCD × F3 med | 0.10 | 0.571 | 0.09 | 0.631 | 0.04 | 0.813 | 0.06 | 0.740 |
| PCD × F3 pos | 1.04 | **<0.001** | 1.04 | **<0.001** | 1.00 | **<0.001** | 1.04 | **<0.001** |
| FC × E | -0.05 | 0.204 | -0.05 | **0.006** | -0.04 | **0.008** | -0.03 | **0.011** |
| E × Pain | 0.00 | 0.633 | 0.00 | **0.030** | 0.01 | **0.001** | 0.01 | **<0.001** |
| FC × E × Target cortex | | | | | | | | |
| FC × E × F3 | -0.23 | 0.095 | -0.16 | **0.008** | -0.16 | **<0.001** | -0.14 | **<0.001** |
| FC × E × F3 ant | 0.04 | 0.787 | -0.06 | 0.386 | -0.08 | 0.073 | -0.06 | 0.126 |
| FC × E × F3 lat | -0.22 | 0.113 | -0.11 | 0.096 | -0.09 | 0.062 | -0.06 | 0.166 |
| FC × E × F3 med | -0.15 | 0.290 | -0.04 | 0.533 | -0.04 | 0.414 | -0.03 | 0.475 |
| FC × E × F3 pos | -0.53 | **<0.001** | -0.20 | **0.005** | -0.09 | 0.072 | -0.02 | 0.640 |
| ICC | 0.13 | | 0.13 | | 0.13 | | 0.13 | |
| N | 15 SubID | | 15 SubID | | 15 SubID | | 15 SubID | |
| Observations | 3502 | | 3502 | | 3502 | | 3502 | |
| Marg. $R^2$ / Cond. $R^2$ | 0.104 / 0.223 | | 0.098 / 0.218 | | 0.101 / 0.219 | | 0.103 / 0.220 | |
| AIC | 12954.810 | | 12975.705 | | 12966.119 | | 12963.635 | |
| AICc | 12955.904 | | 12976.800 | | 12967.214 | | 12964.730 | |

**Notes.** E denotes the shared E-field and target-FC extraction radius, and FC denotes the sgACC seed radius. Columns vary whole-grey-matter FWHM (0, 3, 6 and 12 mm). All models use bilateral sgACC, mean seed time series, median target FC and GSR. Coefficients, ICC and $R^2$ use REML, while information criteria use matched ML fits. P values are nominal.

**Table S10.** Effects of extraction sphere size and smoothing (E = 5 mm, FC = 10 mm).

| | ΔHBC ~ E + FC | | ΔHBC ~ E + FC | | ΔHBC ~ E + FC | | ΔHBC ~ E + FC | |
|---|---|---|---|---|---|---|---|---|
| **ROI E** | **5 mm** | | **5 mm** | | **5 mm** | | **5 mm** | |
| **ROI FC** | **10 mm** | | **10 mm** | | **10 mm** | | **10 mm** | |
| **FC smoothing** | **0 mm** | | **3 mm** | | **6 mm** | | **12 mm** | |
| ***Predictors*** | ***Estimates*** | ***p*** | ***Estimates*** | ***p*** | ***Estimates*** | ***p*** | ***Estimates*** | ***p*** |
| Intercept | 0.25 | 0.160 | 0.26 | 0.149 | 0.27 | 0.140 | 0.29 | 0.107 |
| Session [2] | -0.03 | 0.870 | -0.04 | 0.859 | -0.04 | 0.844 | -0.05 | 0.815 |
| Session [3] | -0.08 | 0.673 | -0.08 | 0.653 | -0.08 | 0.645 | -0.09 | 0.616 |
| Target cortex | | | | | | | | |
| F3 | -0.17 | 0.142 | -0.26 | **0.035** | -0.25 | **0.043** | -0.28 | **0.027** |
| F3 ant | 0.13 | 0.287 | 0.01 | 0.944 | 0.03 | 0.836 | 0.03 | 0.811 |
| F3 lat | 0.06 | 0.649 | -0.01 | 0.950 | 0.06 | 0.641 | 0.09 | 0.501 |
| F3 med | -0.12 | 0.310 | -0.17 | 0.156 | -0.12 | 0.295 | -0.16 | 0.191 |
| F3 pos | -0.24 | **0.038** | -0.27 | **0.021** | -0.23 | **0.040** | -0.22 | 0.068 |
| FC | -2.48 | **<0.001** | -1.41 | **<0.001** | -0.83 | **<0.001** | -0.63 | **0.005** |
| E | 0.01 | **<0.001** | 0.01 | **<0.001** | 0.01 | **<0.001** | 0.01 | **<0.001** |
| Pain | -0.01 | 0.774 | 0.01 | 0.788 | 0.02 | 0.724 | -0.01 | 0.818 |
| Side effects | -0.18 | **<0.001** | -0.18 | **<0.001** | -0.18 | **<0.001** | -0.19 | **<0.001** |
| PCD | 0.02 | 0.742 | 0.02 | 0.709 | 0.01 | 0.839 | 0.01 | 0.765 |
| E × Target cortex | | | | | | | | |
| E × F3 | 0.01 | **0.014** | 0.01 | 0.116 | 0.00 | 0.474 | 0.00 | 0.604 |
| E × F3 ant | 0.00 | 0.924 | -0.00 | 0.735 | -0.00 | 0.468 | -0.00 | 0.587 |
| E × F3 lat | 0.00 | 0.574 | 0.00 | 0.856 | 0.00 | 0.732 | 0.01 | 0.295 |
| E × F3 med | -0.01 | 0.110 | -0.01 | **0.036** | -0.01 | **0.013** | -0.01 | **0.032** |
| E × F3 pos | 0.01 | **0.028** | 0.01 | 0.121 | 0.01 | 0.190 | 0.01 | 0.132 |
| FC × Target cortex | | | | | | | | |
| FC × F3 | 0.58 | 0.801 | -0.70 | 0.545 | -0.75 | 0.352 | -0.09 | 0.903 |
| FC × F3 ant | 3.84 | 0.115 | 0.19 | 0.873 | -0.27 | 0.739 | 0.31 | 0.676 |
| FC × F3 lat | -5.40 | **0.020** | -3.57 | **0.003** | -2.97 | **<0.001** | -2.42 | **0.001** |
| FC × F3 med | 3.25 | 0.125 | 0.83 | 0.440 | 0.05 | 0.943 | 0.42 | 0.523 |
| FC × F3 pos | 5.39 | **0.017** | 1.83 | 0.115 | 1.52 | 0.057 | 1.80 | **0.016** |
| E × PCD | 0.00 | 0.435 | 0.00 | 0.399 | 0.00 | 0.409 | 0.00 | 0.397 |
| PCD × Target cortex | | | | | | | | |
| PCD × F3 | 0.36 | 0.070 | 0.30 | 0.136 | 0.29 | 0.138 | 0.26 | 0.197 |
| PCD × F3 ant | 0.38 | 0.053 | 0.31 | 0.118 | 0.31 | 0.106 | 0.30 | 0.122 |
| PCD × F3 lat | 0.43 | **0.030** | 0.37 | 0.064 | 0.35 | 0.074 | 0.31 | 0.122 |

| | ΔHBC ~ E + FC | | ΔHBC ~ E + FC | | ΔHBC ~ E + FC | | ΔHBC ~ E + FC | |
|---|---|---|---|---|---|---|---|---|
| **ROI E** | **5 mm** | | **5 mm** | | **5 mm** | | **5 mm** | |
| **ROI FC** | **10 mm** | | **10 mm** | | **10 mm** | | **10 mm** | |
| **FC smoothing** | **0 mm** | | **3 mm** | | **6 mm** | | **12 mm** | |
| ***Predictors*** | ***Estimates*** | ***p*** | ***Estimates*** | ***p*** | ***Estimates*** | ***p*** | ***Estimates*** | ***p*** |
| PCD × F3 med | 0.05 | 0.791 | -0.01 | 0.962 | -0.00 | 0.998 | -0.03 | 0.885 |
| PCD × F3 pos | 0.99 | **<0.001** | 0.95 | **<0.001** | 0.97 | **<0.001** | 0.95 | **<0.001** |
| FC × E | -0.11 | **<0.001** | -0.04 | **0.001** | -0.03 | **0.002** | -0.02 | 0.072 |
| E × Pain | 0.00 | 0.086 | 0.00 | **0.032** | 0.00 | **0.009** | 0.00 | **0.006** |
| FC × E × Target cortex | | | | | | | | |
| FC × E × F3 | -0.14 | 0.195 | -0.16 | **<0.001** | -0.14 | **<0.001** | -0.15 | **<0.001** |
| FC × E × F3 ant | -0.15 | 0.192 | -0.12 | **0.021** | -0.08 | **0.015** | -0.10 | **0.001** |
| FC × E × F3 lat | -0.15 | 0.166 | -0.12 | **0.013** | -0.09 | **0.008** | -0.10 | **0.001** |
| FC × E × F3 med | 0.01 | 0.943 | -0.04 | 0.397 | -0.03 | 0.429 | -0.06 | 0.056 |
| FC × E × F3 pos | -0.25 | **0.027** | -0.17 | **0.002** | -0.10 | **0.006** | -0.07 | **0.023** |
| ICC | 0.15 | | 0.15 | | 0.15 | | 0.15 | |
| N | 15 SubID | | 15 SubID | | 15 SubID | | 15 SubID | |
| Observations | 3502 | | 3502 | | 3502 | | 3502 | |
| Marg. $R^2$ / Cond. $R^2$ | 0.105 / 0.240 | | 0.107 / 0.244 | | 0.108 / 0.246 | | 0.107 / 0.240 | |
| AIC | 12945.685 | | 12950.037 | | 12943.251 | | 12940.155 | |
| AICc | 12946.779 | | 12951.132 | | 12944.346 | | 12941.249 | |

**Notes.** E denotes the shared E-field and target-FC extraction radius, and FC denotes the sgACC seed radius. Columns vary whole-grey-matter FWHM (0, 3, 6 and 12 mm). All models use bilateral sgACC, mean seed time series, median target FC and GSR. Coefficients, ICC and $R^2$ use REML, while information criteria use matched ML fits. P values are nominal.

**Table S11.** Effects of extraction sphere size and smoothing (E = 10 mm, FC = 5 mm).

| | ΔHBC ~ E + FC | | ΔHBC ~ E + FC | | ΔHBC ~ E + FC | | ΔHBC ~ E + FC | |
|---|---|---|---|---|---|---|---|---|
| **ROI E** | **10 mm** | | **10 mm** | | **10 mm** | | **10 mm** | |
| **ROI FC** | **5 mm** | | **5 mm** | | **5 mm** | | **5 mm** | |
| **FC smoothing** | **0 mm** | | **3 mm** | | **6 mm** | | **12 mm** | |
| ***Predictors*** | ***Estimates*** | ***p*** | ***Estimates*** | ***p*** | ***Estimates*** | ***p*** | ***Estimates*** | ***p*** |
| Intercept | 0.28 | 0.079 | 0.25 | 0.102 | 0.25 | 0.103 | 0.26 | 0.081 |
| Session [2] | -0.05 | 0.790 | -0.05 | 0.793 | -0.05 | 0.793 | -0.05 | 0.787 |
| Session [3] | -0.10 | 0.585 | -0.10 | 0.591 | -0.10 | 0.591 | -0.10 | 0.588 |
| Target cortex | | | | | | | | |
| F3 | 0.07 | 0.616 | 0.13 | 0.315 | 0.15 | 0.232 | 0.21 | 0.093 |
| F3 ant | 0.31 | **0.017** | 0.39 | **0.002** | 0.39 | **0.002** | 0.44 | **<0.001** |
| F3 lat | 0.41 | **0.001** | 0.48 | **<0.001** | 0.49 | **<0.001** | 0.51 | **<0.001** |
| F3 med | -0.02 | 0.850 | 0.04 | 0.758 | 0.06 | 0.597 | 0.07 | 0.541 |
| F3 pos | 0.02 | 0.867 | 0.10 | 0.404 | 0.11 | 0.352 | 0.13 | 0.269 |
| FC | 0.84 | 0.465 | 1.09 | **0.046** | 0.80 | **0.023** | 0.85 | **0.003** |
| E | 0.01 | **<0.001** | 0.01 | **<0.001** | 0.01 | **<0.001** | 0.01 | **<0.001** |
| Pain | -0.07 | 0.213 | -0.06 | 0.244 | -0.05 | 0.364 | -0.06 | 0.225 |
| Side effects | -0.21 | **<0.001** | -0.21 | **<0.001** | -0.21 | **<0.001** | -0.21 | **<0.001** |
| PCD | 0.01 | 0.901 | 0.00 | 0.955 | 0.01 | 0.879 | 0.01 | 0.866 |
| E × Target cortex | | | | | | | | |
| E × F3 | -0.00 | 0.650 | -0.00 | 0.482 | -0.00 | 0.524 | -0.01 | 0.268 |
| E × F3 ant | -0.01 | 0.137 | -0.01 | 0.141 | -0.01 | 0.099 | -0.01 | 0.061 |
| E × F3 lat | 0.00 | 0.524 | 0.01 | 0.309 | 0.01 | 0.242 | 0.01 | 0.253 |
| E × F3 med | -0.02 | **0.006** | -0.01 | **0.013** | -0.01 | **0.022** | -0.01 | **0.020** |
| E × F3 pos | 0.00 | 0.711 | 0.00 | 0.681 | 0.00 | 0.634 | 0.00 | 0.908 |
| FC × Target cortex | | | | | | | | |
| FC × F3 | 3.63 | 0.417 | -2.42 | 0.224 | -3.04 | **0.011** | -1.76 | 0.056 |
| FC × F3 ant | 5.65 | 0.178 | -1.02 | 0.589 | -2.60 | **0.026** | -1.70 | 0.073 |
| FC × F3 lat | -4.81 | 0.289 | -5.95 | **0.002** | -5.56 | **<0.001** | -4.39 | **<0.001** |
| FC × F3 med | 6.59 | 0.121 | 0.51 | 0.787 | -1.03 | 0.365 | -0.93 | 0.277 |
| FC × F3 pos | 9.45 | **0.023** | 0.82 | 0.658 | -1.08 | 0.344 | -1.32 | 0.143 |
| E × PCD | 0.00 | 0.169 | 0.00 | 0.183 | 0.00 | 0.178 | 0.00 | 0.200 |
| PCD × Target cortex | | | | | | | | |
| PCD × F3 | 0.50 | **0.012** | 0.51 | **0.009** | 0.47 | **0.015** | 0.49 | **0.011** |
| PCD × F3 ant | 0.52 | **0.007** | 0.57 | **0.003** | 0.53 | **0.005** | 0.55 | **0.004** |
| PCD × F3 lat | 0.44 | **0.026** | 0.47 | **0.017** | 0.43 | **0.029** | 0.44 | **0.024** |

| | ΔHBC ~ E + FC | | ΔHBC ~ E + FC | | ΔHBC ~ E + FC | | ΔHBC ~ E + FC | |
|---|---|---|---|---|---|---|---|---|
| **ROI E** | **10 mm** | | **10 mm** | | **10 mm** | | **10 mm** | |
| **ROI FC** | **5 mm** | | **5 mm** | | **5 mm** | | **5 mm** | |
| **FC smoothing** | **0 mm** | | **3 mm** | | **6 mm** | | **12 mm** | |
| ***Predictors*** | ***Estimates*** | ***p*** | ***Estimates*** | ***p*** | ***Estimates*** | ***p*** | ***Estimates*** | ***p*** |
| PCD × F3 med | 0.17 | 0.352 | 0.21 | 0.245 | 0.17 | 0.352 | 0.18 | 0.329 |
| PCD × F3 pos | 1.09 | **<0.001** | 1.14 | **<0.001** | 1.11 | **<0.001** | 1.14 | **<0.001** |
| FC × E | -0.10 | **0.043** | -0.04 | 0.072 | -0.03 | 0.057 | -0.03 | **0.027** |
| E × Pain | 0.00 | **0.015** | 0.00 | **0.003** | 0.01 | **<0.001** | 0.01 | **<0.001** |
| FC × E × Target cortex | | | | | | | | |
| FC × E × F3 | 0.15 | 0.456 | -0.12 | 0.221 | -0.14 | **0.014** | -0.11 | **0.006** |
| FC × E × F3 ant | 0.47 | **0.018** | 0.09 | 0.337 | -0.02 | 0.736 | -0.01 | 0.747 |
| FC × E × F3 lat | 0.34 | 0.097 | 0.04 | 0.696 | -0.04 | 0.472 | -0.03 | 0.546 |
| FC × E × F3 med | 0.57 | **0.007** | 0.17 | 0.092 | 0.05 | 0.422 | 0.03 | 0.454 |
| FC × E × F3 pos | 0.52 | **0.009** | 0.16 | 0.079 | 0.03 | 0.563 | 0.02 | 0.557 |
| ICC | 0.14 | | 0.13 | | 0.13 | | 0.13 | |
| N | 15 SubID | | 15 SubID | | 15 SubID | | 15 SubID | |
| Observations | 3502 | | 3502 | | 3502 | | 3502 | |
| Marg. $R^2$ / Cond. $R^2$ | 0.099 / 0.222 | | 0.103 / 0.222 | | 0.105 / 0.219 | | 0.107 / 0.222 | |
| AIC | 12981.386 | | 12972.006 | | 12961.832 | | 12955.159 | |
| AICc | 12982.480 | | 12973.101 | | 12962.926 | | 12956.253 | |

**Notes.** E denotes the shared E-field and target-FC extraction radius, and FC denotes the sgACC seed radius. Columns vary whole-grey-matter FWHM (0, 3, 6 and 12 mm). All models use bilateral sgACC, mean seed time series, median target FC and GSR. Coefficients, ICC and $R^2$ use REML, while information criteria use matched ML fits. P values are nominal.

**Table S12.** Effects of extraction sphere size and smoothing (E = 10 mm, FC = 10 mm).

| | ΔHBC ~ E + FC | | ΔHBC ~ E + FC | | ΔHBC ~ E + FC | | ΔHBC ~ E + FC | |
|---|---|---|---|---|---|---|---|---|
| **ROI E** | **10 mm** | | **10 mm** | | **10 mm** | | **10 mm** | |
| **ROI FC** | **10 mm** | | **10 mm** | | **10 mm** | | **10 mm** | |
| **FC smoothing** | **0 mm** | | **3 mm** | | **6 mm** | | **12 mm** | |
| ***Predictors*** | ***Estimates*** | ***p*** | ***Estimates*** | ***p*** | ***Estimates*** | ***p*** | ***Estimates*** | ***p*** |
| Intercept | 0.29 | 0.103 | 0.28 | 0.114 | 0.28 | 0.113 | 0.28 | 0.104 |
| Session [2] | -0.05 | 0.808 | -0.05 | 0.810 | -0.05 | 0.806 | -0.05 | 0.789 |
| Session [3] | -0.09 | 0.604 | -0.09 | 0.606 | -0.09 | 0.605 | -0.10 | 0.591 |
| Target cortex | | | | | | | | |
| F3 | -0.12 | 0.330 | -0.16 | 0.201 | -0.16 | 0.201 | -0.12 | 0.347 |
| F3 ant | 0.19 | 0.130 | 0.16 | 0.217 | 0.14 | 0.280 | 0.18 | 0.163 |
| F3 lat | 0.20 | 0.111 | 0.16 | 0.224 | 0.17 | 0.193 | 0.20 | 0.111 |
| F3 med | -0.03 | 0.799 | -0.03 | 0.787 | -0.03 | 0.799 | -0.05 | 0.700 |
| F3 pos | -0.08 | 0.473 | -0.10 | 0.417 | -0.09 | 0.418 | -0.07 | 0.568 |
| FC | -2.09 | **0.020** | -1.14 | **0.010** | -0.77 | **0.008** | -0.30 | 0.216 |
| E | 0.01 | **<0.001** | 0.01 | **<0.001** | 0.01 | **<0.001** | 0.01 | **<0.001** |
| Pain | -0.01 | 0.804 | -0.01 | 0.846 | -0.00 | 0.927 | -0.02 | 0.652 |
| Side effects | -0.22 | **<0.001** | -0.21 | **<0.001** | -0.21 | **<0.001** | -0.21 | **<0.001** |
| PCD | 0.01 | 0.872 | 0.01 | 0.909 | 0.00 | 0.980 | 0.01 | 0.886 |
| E × Target cortex | | | | | | | | |
| E × F3 | 0.00 | 0.569 | 0.00 | 0.917 | 0.00 | 0.947 | -0.00 | 0.998 |
| E × F3 ant | -0.00 | 0.497 | -0.01 | 0.324 | -0.01 | 0.190 | -0.01 | 0.134 |
| E × F3 lat | 0.01 | 0.160 | 0.01 | 0.229 | 0.01 | 0.283 | 0.01 | 0.319 |
| E × F3 med | -0.01 | 0.070 | -0.01 | **0.023** | -0.01 | **0.011** | -0.01 | **0.013** |
| E × F3 pos | 0.01 | 0.128 | 0.01 | 0.244 | 0.00 | 0.395 | 0.00 | 0.588 |
| FC × Target cortex | | | | | | | | |
| FC × F3 | 2.40 | 0.462 | 0.01 | 0.993 | -0.48 | 0.611 | -0.55 | 0.473 |
| FC × F3 ant | 6.43 | **0.045** | 1.70 | 0.272 | 0.22 | 0.820 | 0.06 | 0.939 |
| FC × F3 lat | -7.26 | **0.024** | -4.87 | **0.002** | -3.77 | **<0.001** | -3.12 | **<0.001** |
| FC × F3 med | 2.23 | 0.440 | 0.16 | 0.910 | -0.27 | 0.759 | -0.28 | 0.691 |
| FC × F3 pos | 7.30 | **0.021** | 2.81 | 0.069 | 1.30 | 0.174 | 0.82 | 0.307 |
| E × PCD | 0.00 | 0.400 | 0.00 | 0.324 | 0.00 | 0.299 | 0.00 | 0.264 |
| PCD × Target cortex | | | | | | | | |
| PCD × F3 | 0.43 | **0.029** | 0.41 | **0.038** | 0.41 | **0.038** | 0.40 | **0.042** |
| PCD × F3 ant | 0.48 | **0.012** | 0.47 | **0.014** | 0.45 | **0.018** | 0.46 | **0.016** |
| PCD × F3 lat | 0.42 | **0.031** | 0.42 | **0.034** | 0.41 | **0.036** | 0.39 | **0.048** |

| | ΔHBC ~ E + FC | | ΔHBC ~ E + FC | | ΔHBC ~ E + FC | | ΔHBC ~ E + FC | |
|---|---|---|---|---|---|---|---|---|
| **ROI E** | **10 mm** | | **10 mm** | | **10 mm** | | **10 mm** | |
| **ROI FC** | **10 mm** | | **10 mm** | | **10 mm** | | **10 mm** | |
| **FC smoothing** | **0 mm** | | **3 mm** | | **6 mm** | | **12 mm** | |
| ***Predictors*** | ***Estimates*** | ***p*** | ***Estimates*** | ***p*** | ***Estimates*** | ***p*** | ***Estimates*** | ***p*** |
| PCD × F3 med | 0.09 | 0.623 | 0.09 | 0.636 | 0.09 | 0.622 | 0.09 | 0.633 |
| PCD × F3 pos | 1.04 | **<0.001** | 1.04 | **<0.001** | 1.05 | **<0.001** | 1.05 | **<0.001** |
| FC × E | -0.05 | 0.182 | -0.02 | 0.181 | -0.02 | 0.108 | -0.01 | 0.181 |
| E × Pain | 0.00 | **0.008** | 0.00 | **0.006** | 0.00 | **0.004** | 0.00 | **0.005** |
| FC × E × Target cortex | | | | | | | | |
| FC × E × F3 | -0.40 | **0.009** | -0.23 | **0.002** | -0.14 | **0.002** | -0.11 | **0.002** |
| FC × E × F3 ant | -0.22 | 0.153 | -0.11 | 0.128 | -0.07 | 0.087 | -0.06 | 0.070 |
| FC × E × F3 lat | -0.25 | 0.095 | -0.15 | **0.043** | -0.09 | 0.052 | -0.07 | 0.058 |
| FC × E × F3 med | -0.02 | 0.916 | 0.01 | 0.934 | 0.01 | 0.830 | -0.01 | 0.789 |
| FC × E × F3 pos | -0.09 | 0.539 | -0.04 | 0.564 | -0.02 | 0.625 | -0.01 | 0.722 |
| ICC | 0.15 | | 0.15 | | 0.15 | | 0.14 | |
| N | 15 SubID | | 15 SubID | | 15 SubID | | 15 SubID | |
| Observations | 3502 | | 3502 | | 3502 | | 3502 | |
| Marg. $R^2$ / Cond. $R^2$ | 0.103 / 0.235 | | 0.104 / 0.236 | | 0.105 / 0.235 | | 0.103 / 0.230 | |
| AIC | 12955.553 | | 12947.729 | | 12946.052 | | 12951.231 | |
| AICc | 12956.647 | | 12948.823 | | 12947.146 | | 12952.325 | |

**Notes.** E denotes the shared E-field and target-FC extraction radius, and FC denotes the sgACC seed radius. Columns vary whole-grey-matter FWHM (0, 3, 6 and 12 mm). All models use bilateral sgACC, mean seed time series, median target FC and GSR. Coefficients, ICC and $R^2$ use REML, while information criteria use matched ML fits. P values are nominal.

#### Linear vs. Non-linear models

**Table S13.** Linear vs nonlinear models.

| | **LMM** | | | | **GAM** | | | |
|---|---|---|---|---|---|---|---|---|
| | **ΔHBC ~ E + FC TMS condition == target cortex** | | **ΔHBC ~ E + FC all data** | | **ΔHBC ~ E + FC TMS condition == target cortex** | | **ΔHBC ~ E + FC all data** | |
| ***Predictors*** | ***Estimates*** | ***p*** | ***Estimates*** | ***p*** | ***Estimates*** | ***p*** | ***Estimates*** | ***p*** |
| Intercept | 0.28 | 0.114 | 0.27 | — | 0.52 | **0.019** | 0.37 | — |
| Session [2] | -0.05 | 0.810 | -0.02 | — | -0.06 | 0.756 | -0.02 | — |
| Session [3] | -0.09 | 0.606 | -0.10 | — | -0.10 | 0.631 | -0.10 | — |
| Target cortex | | | | | | | | |
| F3 | -0.16 | 0.201 | -0.08 | — | -0.20 | 0.498 | -0.22 | — |
| F3 ant | 0.16 | 0.217 | 0.03 | — | -0.62 | 0.375 | -0.06 | — |
| F3 lat | 0.16 | 0.224 | -0.02 | — | 0.40 | 0.132 | -0.09 | — |
| F3 med | -0.03 | 0.787 | 0.00 | — | 0.22 | 0.493 | 0.00 | — |
| F3 pos | -0.10 | 0.417 | -0.02 | — | 0.44 | 0.209 | -0.01 | — |
| FC | -1.14 | **0.010** | -0.10 | — | | **<0.001** | | — |
| E | 0.01 | **<0.001** | 0.01 | — | | 0.999 | | — |
| Pain | -0.01 | 0.846 | 0.10 | — | | **<0.001** | | — |
| Side effects | -0.21 | **<0.001** | -0.16 | — | -0.10 | **<0.001** | -0.06 | — |
| PCD | 0.01 | 0.909 | 0.08 | — | | 0.998 | | — |
| E × Target cortex | | | | | | | | |
| E × F3 | 0.00 | 0.917 | 0.00 | — | | 0.695 | | — |
| E × F3 ant | -0.01 | 0.324 | 0.00 | — | | 0.135 | | — |
| E × F3 lat | 0.01 | 0.229 | 0.00 | — | | **0.005** | | — |
| E × F3 med | -0.01 | **0.023** | 0.00 | — | | 0.726 | | — |
| E × F3 pos | 0.01 | 0.244 | 0.00 | — | | **<0.001** | | — |
| FC × Target cortex | | | | | | | | |
| FC × F3 | 0.01 | 0.993 | 0.10 | — | | 0.431 | | — |
| FC × F3 ant | 1.70 | 0.272 | -0.10 | — | | 0.354 | | — |
| FC × F3 lat | -4.87 | **0.002** | 0.19 | — | | 0.471 | | — |
| FC × F3 med | 0.16 | 0.910 | 0.04 | — | | 0.778 | | — |
| FC × F3 pos | 2.81 | 0.069 | 0.22 | — | | 0.578 | | — |
| E × PCD | 0.00 | 0.324 | 0.00 | — | | 0.396 | | — |
| PCD × Target cortex | | | | | | | | |
| PCD × F3 | 0.41 | **0.038** | -0.06 | — | | 0.359 | | — |

| | LMM | | | | GAM | | | |
|---|---|---|---|---|---|---|---|---|
| | **ΔHBC ~ E + FC TMS condition == target cortex** | | **ΔHBC ~ E + FC all data** | | **ΔHBC ~ E + FC TMS condition == target cortex** | | **ΔHBC ~ E + FC all data** | |
| ***Predictors*** | ***Estimates*** | ***p*** | ***Estimates*** | ***p*** | ***Estimates*** | ***p*** | ***Estimates*** | ***p*** |
| PCD × F3 ant | 0.47 | **0.014** | -0.02 | — | | 0.732 | | — |
| PCD × F3 lat | 0.42 | **0.034** | -0.06 | — | | 0.464 | | — |
| PCD × F3 med | 0.09 | 0.636 | -0.00 | — | | **<0.001** | | — |
| PCD × F3 pos | 1.04 | **<0.001** | -0.02 | — | | **<0.001** | | — |
| FC × E | -0.02 | 0.181 | -0.01 | — | | 0.998 | | — |
| E × Pain | 0.00 | **0.006** | 0.00 | — | | **0.031** | | — |
| FC × E × Target cortex | | | | | | | | |
| FC × E × F3 | -0.23 | **0.002** | -0.02 | — | | **<0.001** | | — |
| FC × E × F3 ant | -0.11 | 0.128 | -0.06 | — | | 0.611 | | — |
| FC × E × F3 lat | -0.15 | **0.043** | -0.06 | — | | 0.266 | | — |
| FC × E × F3 med | 0.01 | 0.934 | -0.01 | — | | **0.043** | | — |
| FC × E × F3 pos | -0.04 | 0.564 | -0.02 | — | | **0.003** | | — |
| 5 cm × E | | | | | | 0.312 | | — |
| 5 cm × FC | | | | | | 0.792 | | — |
| 5 cm × E × FC | | | | | | 0.370 | | — |
| 5 cm × PCD | | | | | | 0.690 | | — |
| SubID × Session 1 | | | | | | **<0.001** | | — |
| SubID × Session 2 | | | | | | **<0.001** | | — |
| SubID × Session 3 | | | | | | **<0.001** | | — |
| SubID × E | | | | | | **0.014** | | — |
| ICC | 0.15 | | 0.11 | | — | | — | |
| N | 15 SubID | | 15 SubID | | 15 SubID | | 15 SubID | |
| Observations | 3502 | | 21012 | | 3502 | | 21012 | |

***Notes.*** Columns compare the full LMMs and GAMs in the stimulated-target dataset and the descriptive six-extraction-site extension. Stimulated-target LMM coefficients use REML, six-site LMM coefficients use ML, and GAMs use fREML. Stimulated-target LMM columns show nominal coefficient P values; GAM smooth rows show approximate nominal P values and parametric rows retain coefficients. These P values are unadjusted. Six-site P entries are not reported (—). Descriptive six-site prediction curves are shown in Fig. S2.

#### Six-location prediction curves

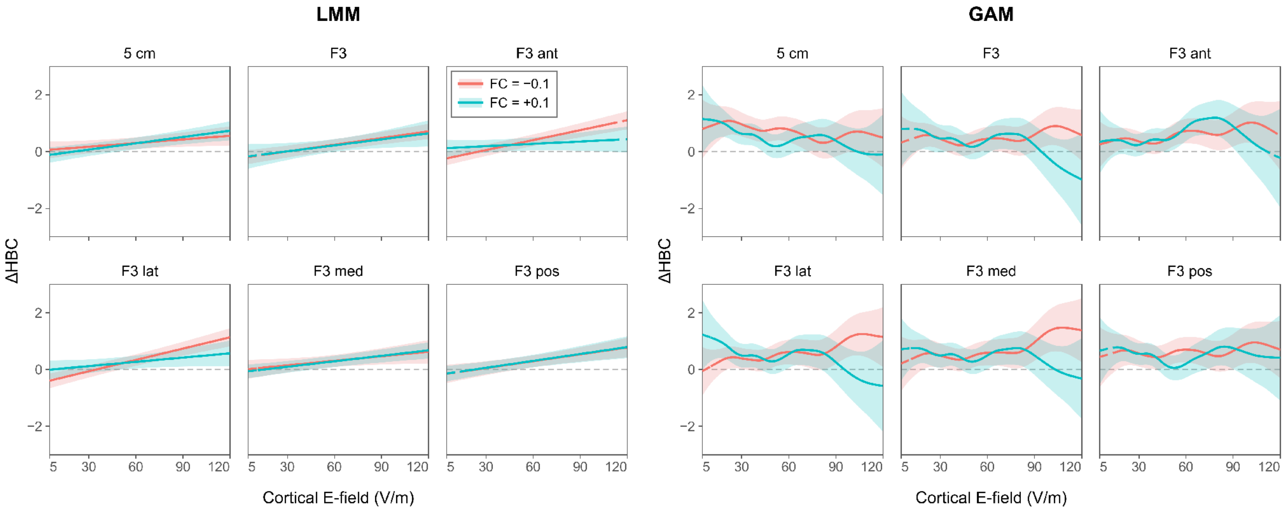


**Figure S2. Descriptive prediction curves from the six-extraction-site extension.** Left: full LMM; right: full GAM. Each recorded stimulation response is represented at six extraction locations (21,012 rows from 15 participants). Both models use the bilateral sgACC seed, mean seed time series, median target FC, 10-mm seed and target radii, 3-mm whole-GM FWHM and GSR. Red and cyan curves represent FC = −0.1 and +0.1, respectively, over E = 5–120 V/m. Shaded bands show model-based 95% intervals. Dashed segments are outside the site-specific observed marginal E or FC range. $\Delta HBC = \log(HBC_{intX}) - \log(HBC_{int0})$.

#### Voxel-wise mapping of HBC modulation

For each participant, raw HBC was regressed on local E-field strength at each native DLPFC cortical element, yielding $R^2$ maps (Fig. 4). The mean of the 15 participant-specific maximum $R^2$ values was 0.136 (SD 0.104). Spatial Pearson correlations between $R^2$ and sgACC FC-z were calculated separately for each participant on paired valid elements, without thresholds on E-field strength, $R^2$, FC sign or significance. Fisher-transformed correlations were tested against zero across 15 participants: $t(14) = -2.242$, P = 0.04165 (two-sided, unadjusted). The FC maps used the bilateral sgACC seed (10-mm radius), unsmoothed mean seed time series, GSR and 8-mm whole-GM FWHM. For display only, negative FC values are divided by the absolute participant-specific minimum FC, and $R^2$ by the participant-specific maximum, on the same paired valid support. The two-colour scales use the same palette in opposite directions; statistical correlations retain all signed, unnormalized FC values.

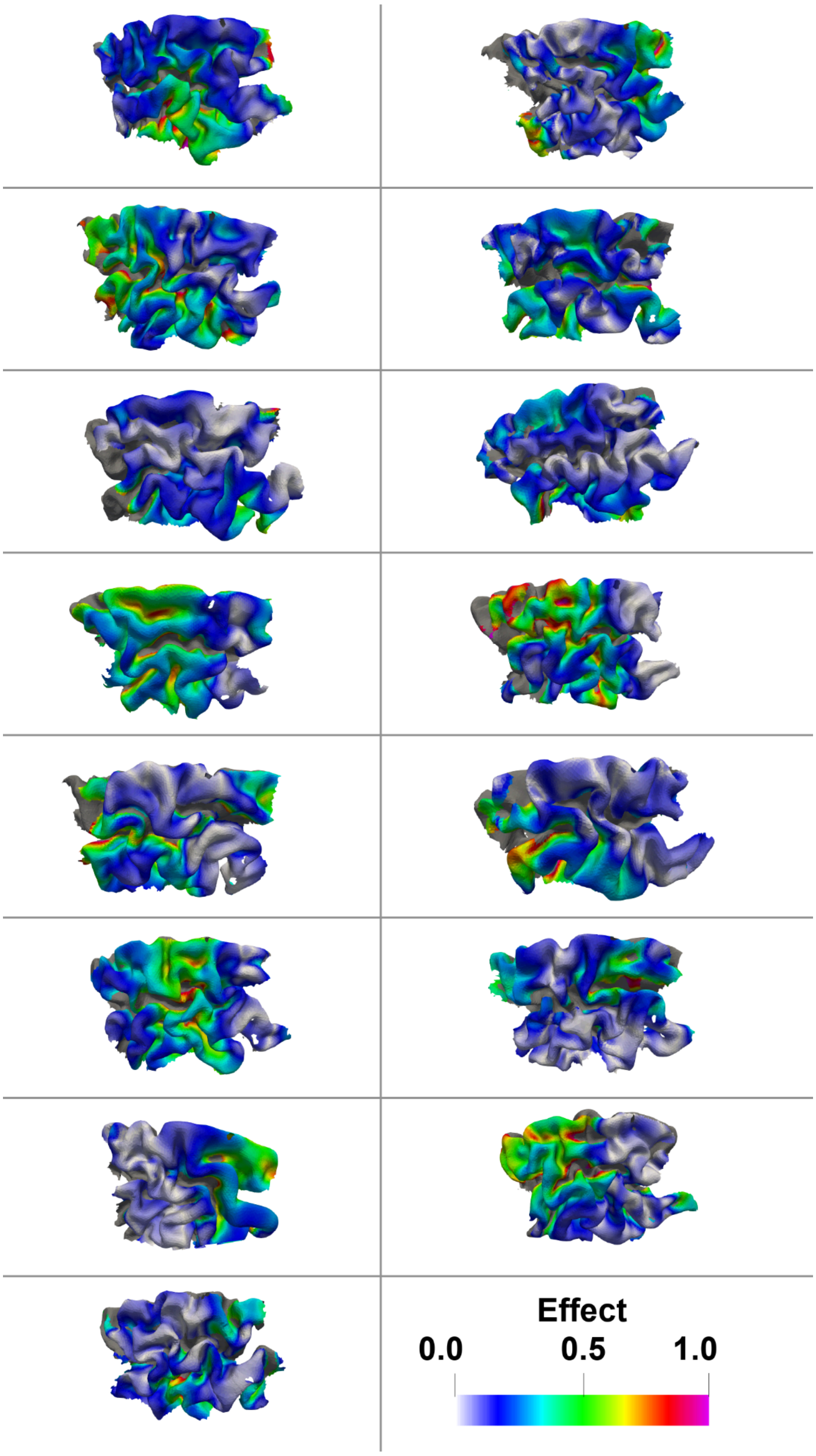


**Figure S3. Individual dose–response gain maps in the DLPFC.** For each participant (cells, same order as Fig. 4), the absolute regression slope of the element-wise linear model relating HBC to local E-field strength was computed across all recorded coil placements, quantifying the local stimulation–response gain (dHBC/dE). Values are normalized by the participant-specific maximum gain (colour scale "Effect", 0–1), so that warm colours indicate cortical locations where variation in E-field strength is translated most strongly into HBC. Cortical elements with local E-field variance below the participant-specific 25th percentile are masked (grey), as slope estimates are unstable where the E-field varies little across stimulation conditions. The gain maps closely mirror the explained-variance ($R^2$) maps in Figure 4, correlated at $r = 0.84 \pm 0.03$ (mean ± SD) across participants, confirming that regions of high E-field–HBC predictability coincide with regions of strong dose–response coupling.
